# rDNA breaks activate dsRNA pattern recognition through sense-antisense transcription

**DOI:** 10.64898/2026.05.26.728043

**Authors:** Jie Chen, Sangin Kim, Sixing Chen, Zongchao Mo, Yiyang Jiang, Kathy Fange Liu, Qin Li, Roger A. Greenberg

**Affiliations:** Department of Cancer Biology, Penn Center for Genome Integrity, Basser Center for BRCA, Perelman School of Medicine, University of Pennsylvania, Philadelphia, PA 19104, USA; Department of Genetics, University of Pennsylvania, Philadelphia, PA 19104, USA; Department of Genetics, Penn Center for Genome Integrity, Penn Institute for Immunology and Immune Health (I3H), Perelman School of Medicine, University of Pennsylvania, Philadelphia, PA 19104, USA; Department of Biochemistry and Biophysics, Penn Center for Genome Integrity, Penn Institute for RNA Innovation, Perelman School of Medicine, University of Pennsylvania, Philadelphia, PA 19104, USA

**Author notes:** Corresponding author, Roger A. Greenberg. These authors contributed equally to this work (Jie Chen, Sangin Kim).

## Abstract

Myriad DNA damaging chemo- and radio- therapies interfere with ribosomal RNA (rRNA) transcription and processing, yet the biological consequences of these phenomena remain unclear. Here we show that aberrant transcripts emanating from rDNA breaks engage double-stranded RNA pattern recognition receptors, melanoma differentiation-associated protein 5 (MDA5) and retinoic acid-inducible gene I (RIG-I) to activate immune signaling. rDNA damage abolishes full-length rRNA synthesis while generating truncated sense-antisense rRNA transcripts, whose accumulation is restrained by ATM and ATR kinase activities. Purification of the endogenous MDA5-filament coupled with sequencing identified complementary sense and antisense rRNA transcripts that terminate and initiate near the break site, respectively. This implicates aberrant rRNA species as a major source of damage induced endogenous ligands for dsRNA pattern recognition and establishes a mechanism by which nucleolar stress is coupled to immune signaling.

## Introduction

The DNA damage and innate immune responses each rely on sensors that detect the presence of aberrant nucleic acid structures in the nucleus and cytoplasm, respectively. Myriad genomic lesions are rapidly detected to elicit cognate DNA repair mechanisms (*1, 2*). In an analogous manner, the presence of double-stranded DNA (dsDNA) or double-stranded RNA (dsRNA) in the cytoplasm triggers molecular pattern recognition and signaling to transcription factors that promote interferon-stimulated gene (ISG) expression. Sensing of cytosolic DNA or dsRNA occurs by pattern recognition receptors (PRRs) cyclic GMP-AMP Synthase (cGAS) and melanoma differentiation-associated gene 5 (MDA5) and retinoic acid-inducible gene I (RIG-I), respectively. These PRRs are critical to mount productive innate immune responses to intracellular viruses and bacteria (*3–5*).

Endogenous nucleic acids also engage cytosolic pattern recognition receptors in a process termed viral mimicry. Cellular dsDNA and dsRNA elicit responses similar to those triggered by viral genomes, resulting in interferon-stimulated gene expression and the pediatric type I interferonopathy Aicardi-Goutières Syndrome (*6–11*). The endogenous DNA and dsRNA species that are responsible for excessive pattern recognition remain largely undefined, albeit both mitochondrial and genomic DNA are linked to cGAS activation (*12–16*). Several sources of immunogenic dsRNA have also been reported, including mitochondrial dsRNA, inverted repeat Alu elements in patients with hyperactivating mutations in MDA5, and derepressed endogenous retroviral elements following treatment with agents that inhibit cytosine methylation (*11, 17, 18*). DNA damage has been shown to activate both cGAS and the RNA sensors MDA5 and RIG-I (*12, 14–16, 19–27*). ATM or ATR inhibition combined with DNA damaging agents elicits innate immune signaling predominantly via dsRNA sensing pathways, albeit mechanism underlying this observation remain unclear (*21–23*). This major gap in understanding stems from current inabilities to isolate the active forms of MDA5 and RIG-I, which exist as filaments on their dsRNA ligands (*28–31*).

In this study, we report that rDNA double-strand breaks (DSBs) produce immunogenic sense-antisense dsRNA transcripts that activate MDA5 and RIG-I. ATM and ATR restrain production of both sense and antisense rRNAs at damaged rDNA. Moreover, purified endogenous MDA5 filaments harbored complementary sense and antisense rRNA transcripts that terminate and initiate at the break site, respectively. These findings establish a general mechanism by which DNA damage links nucleolar stress to immune signaling.

### AhTM- and ATR- mediated transcriptional silencing at rDNA breaks restrains innate immune activation

To investigate how the genomic location of DSBs influences dsRNA pattern recognition, we compared innate immune signaling following rDNA damage using the I-PpoI endonuclease versus genome-wide DSB induction by AsiSI. I-PpoI primarily creates DSBs in the rDNA and has approximately ten cut sites in non-rDNA genomic sites (*32*). AsiSI is a methylation sensitive endonuclease that creates DSBs throughout the nucleus with minimal cleavage in the rDNA (*33*). Damage localization by γH2AX staining confirmed the reported break induction by each nuclease (**Fig. 1A**) (*34, 35*). DSB induction was performed in hTERT immortalized retinal pigmented epithelial (RPE-1) cells, which lack cGAS expression but are competent for both MDA5 and RIG-I dsRNA sensing. This allowed us to assess damage induced dsRNA contributions to ISG responses without interference from dsDNA pattern recognition. Although comparable levels of DNA damage occurred in each setting, breaks induced by I-PpoI triggered much higher levels of STAT1 phosphorylation (p-STAT1, Tyr 701) and induction of the interferon-stimulated gene 56 (ISG56) (**Fig. 1B**).

**Figure 1.**
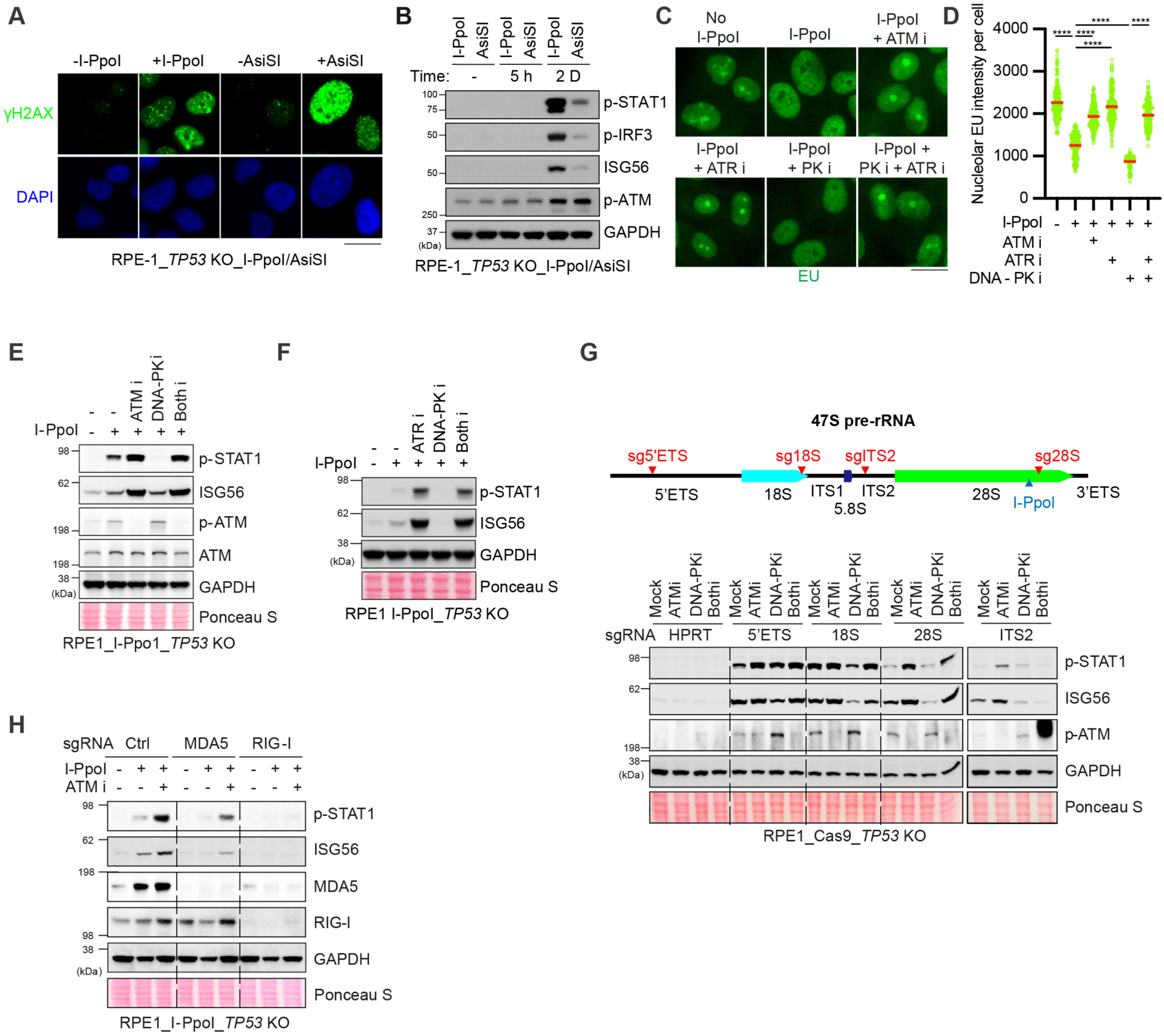
Transcriptional silencing at rDNA breaks limits inflammatory signaling. **(A)** Representative images of γH2AX immunostaining with following I-PpoI- or AsiSI-mediated DNA double-strand breaks. Scale bar, 20 μm. **(B)** Immunoblots confirming DNA damage and immune activation induced by I-PpoI or AsiSI. **(C, D)** Immunofluorescence of nucleolar RNA synthesis visualized by EU-click labeling with representative images and quantification. For EU analysis, at least 150 cells were quantified per experiment. Three independent experiments were performed, and one representative was displayed. Bars represent mean ± SEM. Statistical significance was determined by an unpaired two-tailed Student’s *t*-test. **** p < 0.0001. Scale bar, 20 μm. **(E, F)** Immunoblots of whole-cell lysates showing STAT1 phosphorylation and ISG56 expression 48 h after I-PpoI-mediated rDNA breaks in the presence of ATM, ATR or DNA-PK inhibition in hTERT RPE-1 *p53* KO cells. **(G)** (Top) Schematic of the human rDNA showing the 47S pre-rRNA transcription unit and the positions of sgRNAs targeting the 5’ETS, 18S, ITS2, and 28S regions, along with the I-PpoI recognition site. (Bottom) Immunoblots of whole-cell lysates following transfection of rDNA-targeting sgRNAs in Cas9-expressing hTERT RPE-1 *p53* KO cells. **(H)** Immunoblots of whole-cell lysates assessing the dependence of the cytosolic RNA sensors MDA5 and RIG-I on I-PpoI-induced immune activation in the presence or absence of ATM inhibition.

Given that approximately 50-70% of cellular transcription originates from rDNA repeat arrays within the nucleolus, we hypothesized that damage induced transcriptional perturbation of rRNA may be particularly immunogenic. Consistent with previous reports (*19, 32, 36–40*), induction of rDNA breaks by I-PpoI caused rapid transcriptional silencing within nucleoli as assessed by nascent RNA synthesis using 5’-ethynyl uridine (EU) incorporation (**Fig. 1C, D**). This response required ATM and ATR kinase activity, as inhibition of either kinase attenuated rDNA transcriptional silencing, whereas inhibition or genetic loss of non-homologous end-joining (NHEJ) factors DNA-PKcs, XRCC4, or Ligase 4 led to persistent breaks and further reinforced transcriptional repression (**Fig. 1C, D**, and **Fig. S1A-D**). rDNA transcriptional silencing was tightly linked to innate immune activation. Inhibition of ATM or ATR markedly enhanced STAT1 phosphorylation and ISG56 expression following I-PpoI induction, while DNA-PK inhibition suppressed immune activation in a dose-dependent manner (**Fig. 1E, F**, **Fig. S1B**). To determine if damage across the rDNA locus could elicit ATM and ATR dependent silencing and innate immune signaling, we targeted distinct regions of the rDNA repeat using CRISPR/Cas9 coupled with single guide RNAs (sgRNAs) that included the 5′-external transcribed spacer (5′-ETS), 18S, internal transcribed spacer 2 (ITS2), or 28S rDNA sites. Targeting any of these regions induced rRNA transcriptional silencing and phenocopied I-PpoI-induced immune activation, including ISG responses that were enhanced by ATM inhibition (**Fig. 1G, Fig. S1E, F**). Depletion of either MDA5 or RIG-I markedly suppressed innate immune activation following rDNA breaks and ATM inhibition (**Fig. 1H, Fig. S1G, H**). Notably, RIG-I depletion resulted in stronger suppression of ISG responses and loss of damage-induced increases in MDA5. This data is consistent with RIG-I being a more potent sensor of rDNA damage that triggers a feed forward mechanism of dsRNA pattern recognition by both MDA5 and RIG-I in response to rDNA damage.

### rDNA damage results in nascent rRNA association with MDA5 and RIG-I

To test whether rDNA damage leads to physical association between nascent RNA and PRRs, we performed proximity ligation assays (PLA) between 5′-ethynyl uridine (EU)-labeled nascent RNA that was detected by biotin-azide click chemistry, and MDA5 or RIG-I using specific antibodies (**Fig. 2A, B**). PLA signals were robustly detected in the cytosol and enhanced by ATM or ATR inhibition at one hour after EU pulse in cells expressing I-PpoI, but not AsiSI (**Fig. S2, 3**). PLA signals increased up to 48 hours after damage induction, were unaffected by interferon-β (IFN-β)-mediated upregulation of MDA5 or RIG-I, and were almost completely suppressed in MDA5- or RIG-I deficient cells (**Fig. S4**), confirming the specificity of this assay.

**Figure 2.**
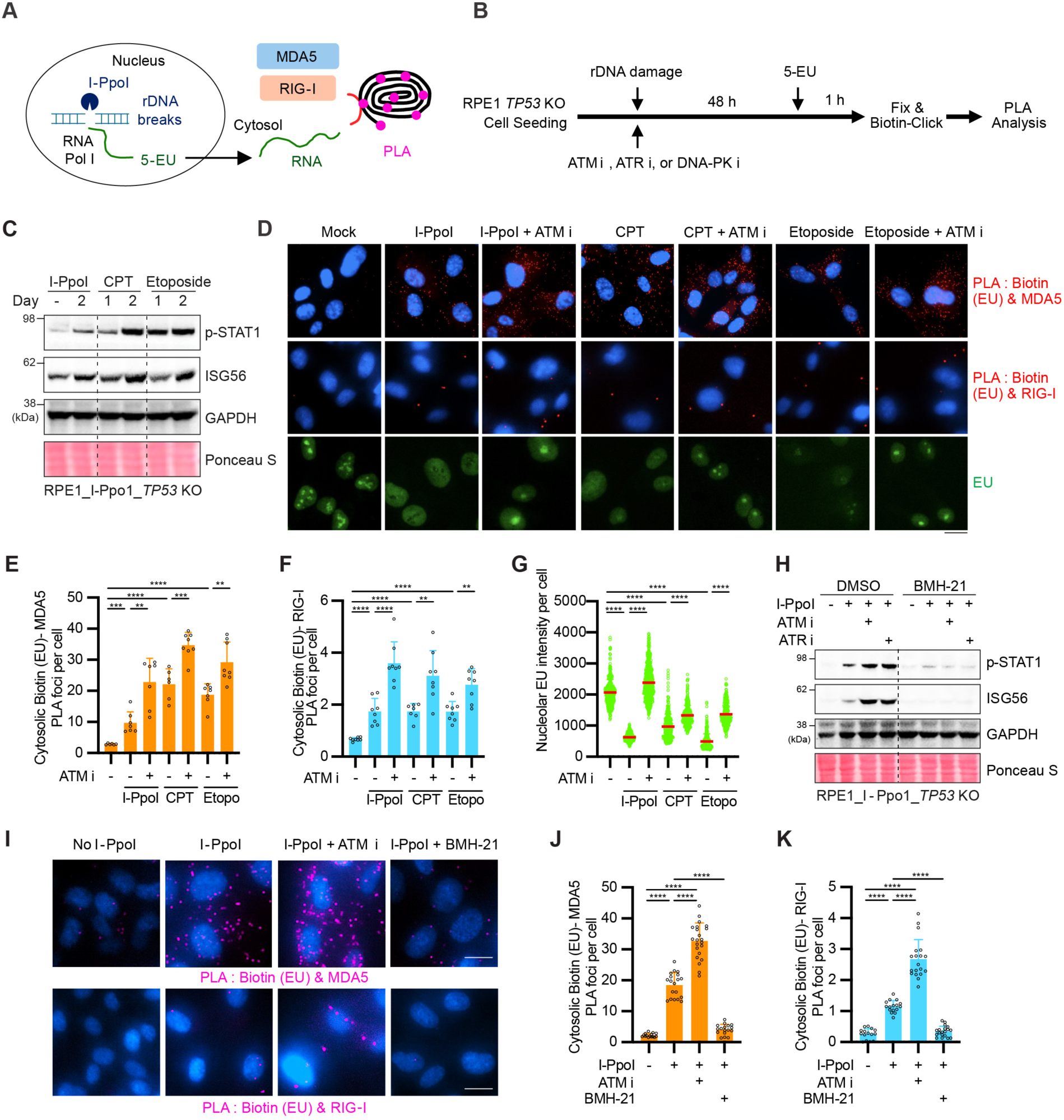
Nascent rRNA associates with MDA5 and RIG-I after damage. **(A)** Schematic diagram of proximity ligation assay (PLA) between biotin-labeled nascent RNA and the cytosolic RNA pattern recognition receptors MDA5 and RIG-I following rDNA breaks. Nascent RNA was labeled with 5’-ethynyl uridine (EU) and biotinylated by click chemistry prior to PLA detection. (**B**) Experimental scheme for detecting cytosolic nascent RNA sensing by MDA5 or RIG-I. **(C)** Immunoblots of p-STAT1 and ISG56 following I-PpoI induction, camptothecin (CPT), or etoposide treatment. **(D)** Representative images of PLA following EU labeling and biotin click reaction after DNA damage. (Top) PLA signals indicate proximity between biotin-labeled nascent RNA (EU) and MDA5 or RIG-I. (Bottom) Nucleolar RNA synthesis visualized by EU-click labeling. Scale bar, 20 μm. **(E-G)** Quantification of cytosolic PLA signals EU-labeled nascent RNA and MDA5 (E), RIG-I (F), and nucleolar EU signal intensity (**G**). (**H**) Immunoblots of p-STAT1 and ISG56 following I-PpoI-mediated rDNA breaks in the presence or absence of RNA polymerase I inhibitor BMH-21. **(I)** Representative images of cytosolic PLA foci between biotin-labeled nascent RNA (EU) and MDA5 or RIG-I following BMH-21 treatment. Scale bar, 20 μm. **(J, K)** Quantification of cytosolic PLA signals between biotinylated EU-labeled nascent RNA and MDA5 (J) or RIG-I (K). (**C-K**) Experiments were performed in hTERT RPE-1 *p53* KO cells with inducible expression of I-PpoI. (**E, F, J, K**) Each dot represents the mean value of at least 50 cells. (**G**) For EU analysis, at least 150 cells were quantified per experiment. Three independent experiments were performed, and one representative result is displayed. (**E-G, J, K**) Bars represent mean ± SEM. Statistical significance was determined by unpaired two-tailed Student’s *t*-test. **** p < 0.0001, *** p <0.001, ** p < 0.01.

To determine whether innate immune activation is a general consequence of rDNA damage, we induced DNA breaks using the clinically relevant topoisomerase inhibitors camptothecin (CPT) or etoposide, which are known to damage the rDNA (*41–43*). Both CPT and etoposide treatments triggered robust STAT1 phosphorylation and ISG56 expression (**Fig. 2C**). Moreover, CPT or etoposide treatment induced rDNA transcriptional silencing and promoted cytosolic PLA signals between nascent RNA and MDA5 or RIG-I. ATM- or ATR-inhibition restored rRNA transcription and enhanced PLA signals between nascent RNA and MDA5 or RIG-I (**Fig. 2D-G**). Irradiation (IR) also induced strong immune activation and cytosolic PLA signals (**Fig. S5A-E)** that were enhanced by ATM or ATR inhibition. We next tested whether RNA sensing requires ongoing RNA polymerase I transcription. Inhibition of RNA polymerase I elongation, using BMH-21 (*44*), markedly reduced rRNA transcription and cytosolic PLA signals between nascent RNA and MDA5 or RIG-I (**Fig. 2H-K, Fig. S5F, G**). Collectively, these results indicate that diverse rDNA damaging modalities encompassing nucleases, chemotherapy, or ionizing radiation generate nascent rRNA species that rapidly associate with MDA5 and RIG-I to activate innate immune signaling. Moreover, ATM- and ATR-mediated rDNA silencing serves to attenuate this response.

### rDNA breaks disrupt pre-rRNA processing and nucleolar structure

We hypothesized that immunogenic dsRNAs are derived from aberrant rRNA species produced in the context of rDNA damage. We therefore examined how rDNA breaks affect processing of nascent 47S pre-rRNA (**Fig. 3A**). rDNA breaks by I-PpoI suppressed intact 45S pre-rRNA and caused accumulation of truncated 45S and 32S pre-rRNA species, with commensurate reduction in the 12S intermediate (**Fig. 3B**). ATM inhibition further increased the abundance of these truncated rRNA species, whereas inhibition of DNA-PK reduced their accumulation. To determine whether disruption at distinct rDNA loci differentially affects rRNA maturation, we targeted individual regions of the rDNA repeat, including the 5′-ETS, 18S, ITS2, or 28S regions. Targeting each region resulted in the accumulation of truncated rRNA species consistent with the expected processing intermediates for the respective cleavage sites (**Fig. 3C, D**). Accumulation of these truncated rRNA species across multiple rDNA loci was accompanied by cytosolic rRNA sensing by MDA5, and RIG-I (**Fig. S6**). These results indicate that disruption at multiple positions across the rDNA array interferes with coordinated 47S pre-rRNA maturation and is associated with immunogenic RNA sensing.

**Figure 3.**
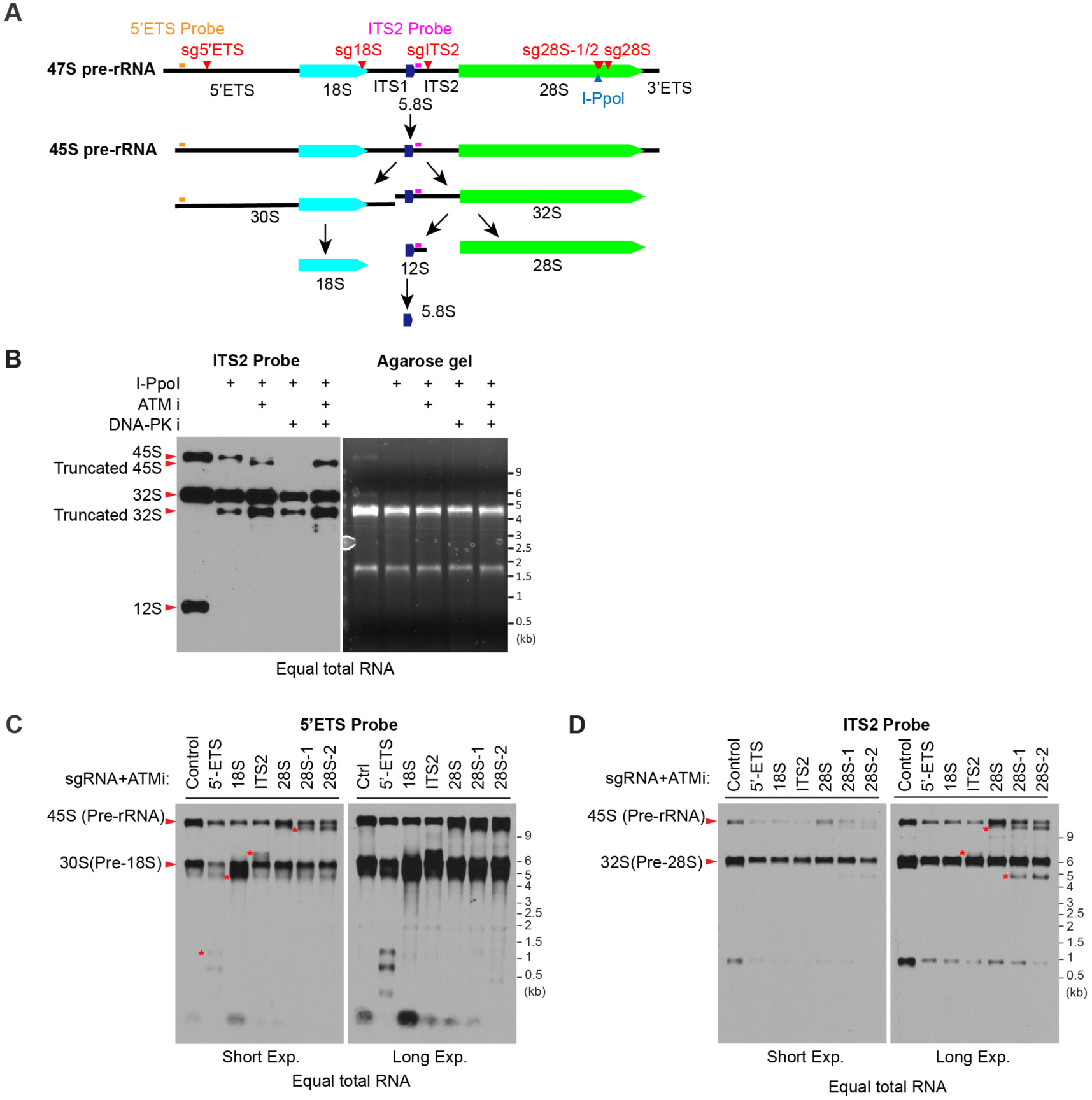
rDNA breaks disrupt nascent rRNA processing. **(A)** Schematic overview of pre-rRNA processing following transcription of the 47S pre-rRNA precursor, indicating the 5′-ETS, 18S, ITS1, ITS2, and 28S regions, the positions of sgRNA targeting sites, the I-PpoI cleavage site, and the locations of the 5’ETS and ITS2 probes. **(B)** Northern blot analysis using an ITS2 probe shows truncated 45S and 32S rRNA pre-rRNA species that are approximately 1.3 kb shorter than their full-length counterparts following I-Ppo1-mediated rDNA breaks, consistent with cleavage within the 28S regions (Left) and an agarose gel analysis of 28S and 18S rRNA from purified total RNA (Right). **(C, D)** Northern blot analysis of truncated rRNA species following sgRNA-mediated rDNA breaks targeting the 5′-ETS, 18S, ITS2, or 28S regions, detected by the 5’-ETS (C) and ITS2 (D) probes, respectively. (**C**) The 5’ETS probe showed that sgRNAs targeting the 5′-ETS, 18S, ITS2, or 28S regions induced truncated pre-rRNA species of approximately 1.2kb, 5.1kb, 6.7kb and 11.3kb, respectively, as indicated by asterisk. (**D**) The ITS2 probe showed that sgITS2 induced a truncated 6.7 kb pre-rRNA and sg28S induced a truncated 11.3 kb pre-rRNA, which were further process into truncated 4.5 kb 32S RNA.

Faithful rRNA biogenesis depends on the higher-order organization of the nucleolus, which comprises fibrillar centers (FCs), the dense fibrillar component (DFC), and the granular component (GC) (*45–48*) (**Fig. S7A**). Accordingly, rDNA damage led to pronounced disruption of nucleolar architecture, as evidenced by altered colocalization and redistribution of FC and DFC markers, including RPA194, UBF, and fibrillarin (FBL) (**Fig. S7B-E**). ATM inhibition resulted in fewer distinct nucleolar structures, a phenotype consistently observed by EU labeling (**Fig. 1C**, **2D and Fig. S1C, 1E, 5F**). Together, rDNA breaks disrupt coordinated rRNA processing and nucleolar organization. These events may contribute to the generation of aberrant rRNA species that are detected by cytosolic dsRNA-sensing pathways.

### Active MDA5 filaments contain rRNA-derived sense and antisense dsRNAs that are produced at rDNA breaks

Purification of endogenous MDA5 filaments from cells has not been reported, thus preventing direct identification of its dsRNA ligands. The E3 ligase TRIM65 specifically recognizes active MDA5 filaments through bivalent interactions but not inactive MDA5 monomers (*49, 50*). We leveraged this information to isolate the active MDA5 filament and identify its associated dsRNA ligands. Purified recombinant Streptavidin-Binding Peptide (SBP)-tagged TRIM65 fragments inclusive of the coiled-coil and PRY-SPRY domains were used to pull down active MDA5 filaments in control and I-PpoI induced RPE-1 cells (**Fig. 4A, B)**. Recombinant TRIM65 copurified with MDA5 but not with RIG-I from cellular lysates (**Fig. 4B**). TRIM65 and MDA5 maintained interaction after high-concentration RNase A digestion (1 mg/ml), which was used to reduce nonspecific RNA association and enhance signal-to-noise ratios (**Fig. 4A, B**).

**Figure 4.**
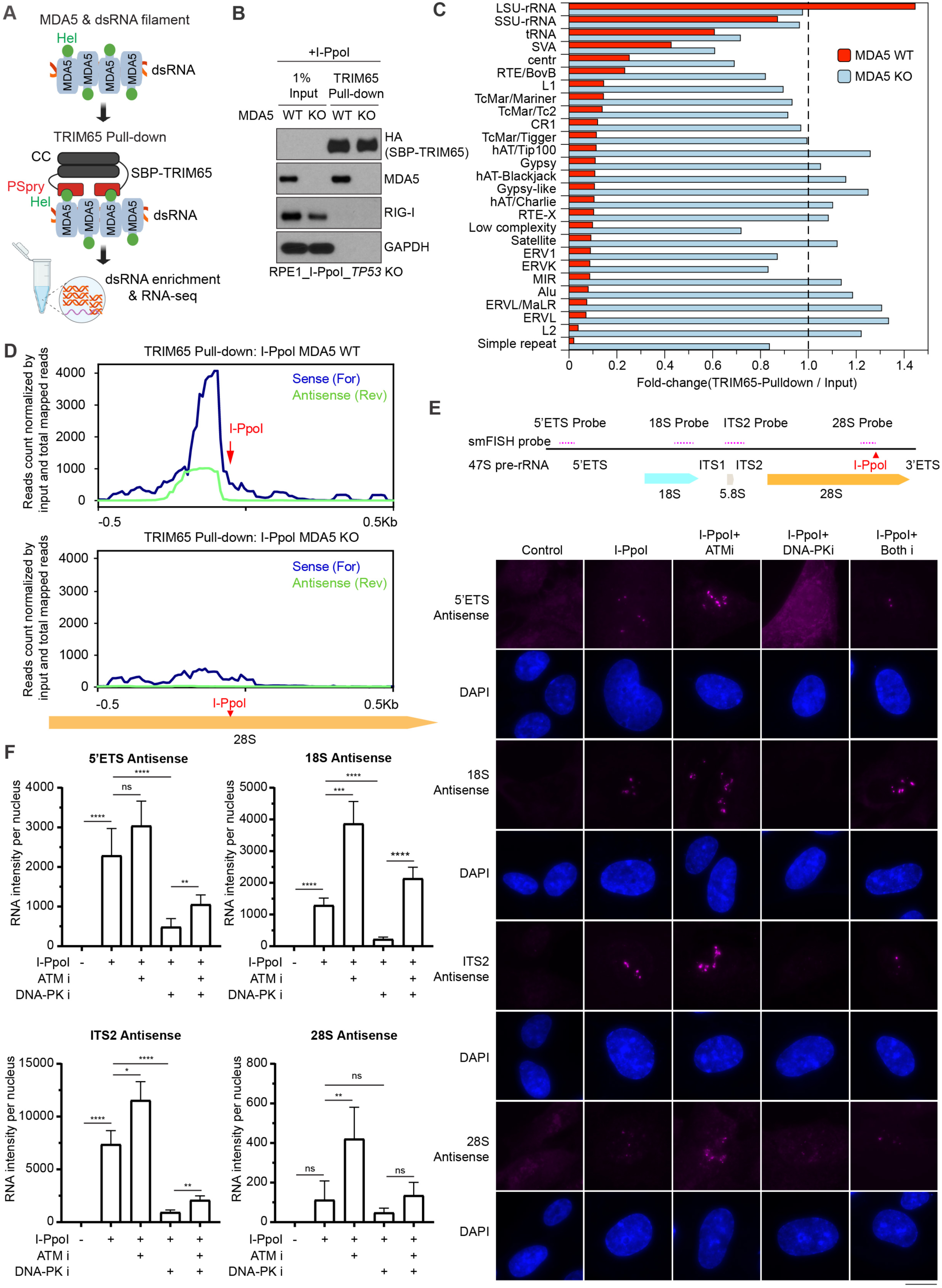
MDA5 associates with rRNA-derived sense and antisense dsRNAs. **(A)** Schematic diagram of the TRIM65-mediated pulldown strategy used to purify MDA5-associated RNA and identify immunogenic rRNA species by RNA sequencing. Hel, Helicase; CC, Coiled-coil; PSpry, Pry-Spry domains; SBP, Streptavidin-binding peptide. **(B)** Immunoblots showing TRIM65 pulldown and co-purification of MDA5 in MDA5 wild-type (WT) but not in knockout (KO) cells. (**C**) Fold enrichment of repetitive elements in TRIM65 pull-downs from the indicated conditions (MDA5 WT and MDA5 KO) normalized to the corresponding input and grouped by repeat class. (**D**) Strand-specific signal of TRIM65-pulldown for sense (forward) and antisense (reverse) strands near the I-PpoI cutting site in MDA5 WT (upper panel) and MDA5 KO (lower panel) cells. Pulldown signal was normalized against the corresponding input for ±500 bp window. (**E**) Top, schematic showing the site-specific locations of probes targeting the 5′-ETS, 18S, ITS2, and 28S regions. Bottom, representative images of smFISH following I-PpoI-mediated rDNA breaks in the presence of ATM or DNA-PK inhibition using antisense 5’ETS, 18S, ITS2, or 28S probes. (**F**) Quantification of antisense RNA intensity per nucleus with corresponding probes. At least 50 cells were quantified per experiment. Three independent experiments were performed, and one representative result is shown. Bars represent mean ± SEM. Statistical significance was determined by an unpaired two-tailed Student’s *t*-test. **** p < 0.0001, *** p <0.001, ** p < 0.01, * p <0.05, ns, not significant. Scale bar, 20 μm.

RNA sequencing of TRIM65 pull-downs following I-PpoI-mediated rDNA break induction revealed an enrichment of specific rRNA-derived sequences compared to TRIM65 pulldowns in MDA5 KO controls (**Fig. 4C**). Reads mapped to the 28S rRNA (a component of the large subunit, LSU) were enriched in MDA5 wild-type cells, whereas other repetitive RNA species were depleted. Notably, Alu repeats, the major endogenous substrate of ADAR1-editing and proposed source of MDA5 ligands in genetic conditions associated with elevated type I interferon (*18, 51*) were not enriched by our TRIM65 pulldown. Similarly, mitochondrial RNA was not enriched in the MDA5 filament purification. This suggests a distinct mechanism of immunogen generation by rDNA breaks. Strand-specific analysis of rRNA-derived reads revealed pronounced enrichment of both forward (sense) and reverse (antisense) strand reads mapping to a ∼200 base region in the vicinity of the I-PpoI cleavage site (**Fig. 4D**), consistent with the formation of rRNA-derived sense-antisense RNA duplexes near rDNA breaks. Neither 28S sense nor antisense transcripts were present in the TRIM65 pulldown from MDA5-deficient cells attesting to the specificity of damage dependent immunogenic dsRNAs being produced in proximity to the rDNA damage site.

Consistent with the detection of MDA5-associated sense and antisense rDNA transcripts, we observed dsRNA accumulation at nucleolar caps following I-PpoI- or sg18S-induced rDNA breaks (**Fig. S8A, B**). This signal was sensitive to RNase A and RNase III, but not RNase H, consistent with the presence of dsRNA. To directly visualize both sense and antisense rRNA transcription following rDNA breaks, we performed strand-specific single-molecule fluorescence *in situ* hybridization (smFISH) with probes complementary to anti-sense 5′-ETS, ITS2, 18S, or 28S transcripts. Antisense transcripts emerged beginning ∼ 4 hours after sgRNA transfection and progressively accumulated over time (**Fig. S8C-F**). They were sensitive to inhibition of RNA polymerase I by BMH-21 and were not detected with control breaks at the HPRT locus (sgCtrl). We further tested dependency on regulators of DSB silencing. I-PpoI break induction produced antisense transcripts at all the regions, including 28S, ITS2, 18S, and 5’ETS, that were quantitatively enhanced by ATM inhibition and reduced by DNA-PK inhibition (**Fig. 4E, F**). Together, these findings support a model in which rDNA damage induces bidirectional rRNA transcription. ATM and ATR dependent DSB silencing serves to limit the immunogenic rRNA-derived sense-antisense dsRNA species that selectively associate with dsRNA sensing PRRs.

### Immunogenic rRNAs are generated upstream of the rDNA break site by sense and antisense transcription

Although transcription is rapidly silenced at damaged rDNA loci in the nucleolus (**Fig. S4F**), it can be re-initiated during or after DNA damage repair. Strand-specific single-molecule fluorescence *in situ* hybridization (smFISH) targeting the 5′ ETS, 18S, ITS2, or 28S regions revealed that rDNA breaks induce position-dependent antisense rRNA transcription (**Fig. 5A**). Antisense 5′ ETS transcripts were detected following break induction at all tested rDNA regions, whereas antisense 18S transcripts were detected after breaks at the 18S, ITS2, or 28S regions, but not after 5′ ETS targeting. Antisense ITS2 transcripts were detected only when breaks were introduced at the ITS2 or 28S regions, but not after 5′ ETS or 18S targeting (**Fig. 5A, B**). These patterns indicate that rDNA damage induces antisense transcription that extends from break-proximal regions toward upstream rDNA sequences. Antisense transcripts were strictly rDNA damage-dependent and undetectable in unperturbed cells, providing direct evidence of rDNA damage-dependent bidirectional rRNA transcription. We noticed that antisense RNA signals were diminished in the vicinity of the corresponding break site for all sgRNAs tested (28S, ITS2, 18S, and 5’ETS). This may suggest variability in antisense RNA transcription initiation or processing. Despite these potential issues, we still observed excellent colocalization of sense-antisense 5’ETS transcripts by smFISH in cells with sg5’ETS transfection but not in sgCtrl cells, indicative of immunogenic dsRNA formation in the vicinity of the break site (**Fig. 5C, D**).

**Figure 5.**
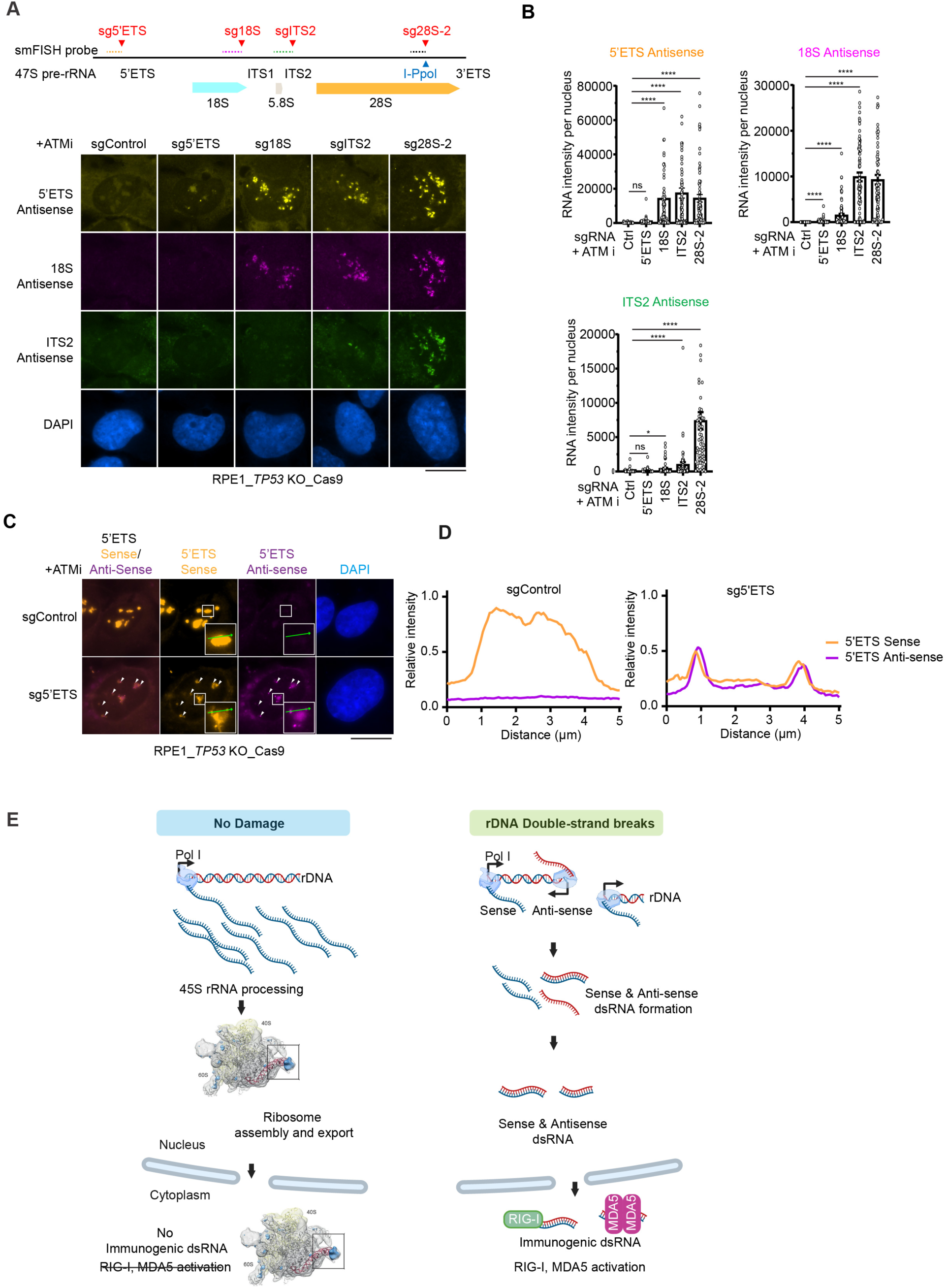
Antisense transcription initiates near the rDNA break-site and extends upstream through the 5’ETS region. (**A**) Top, schematic showing the site-specific locations of probes targeting the 5′-ETS, 18S, ITS2, and 28S regions. Bottom, representative strand-specific smFISH images of nascent RNA following gRNA-mediated targeting of the 5’ ETS, 18S, ITS2, or 28S regions in the presence of ATM inhibition. Antisense transcripts were visualized using strand-specific probes. Scale bar, 20 μm. (**B**) Quantification of antisense smFISH RNA intensity per nucleus following rDNA break using indicated gRNAs in the presence of ATM inhibition. At least 50 cells were quantified per experiment. Three independent experiments were performed, and one representative result is shown. Bars represent mean ± SEM. Statistical significance was determined by an unpaired two-tailed Student’s t-test. **** p < 0.0001, ns, not significant. (**C**) Representative images of sense and antisense smFISH RNA following gRNA-mediated rDNA break generation targeting 5’ ETS in the presence of ATM inhibition. Scale bar, 20 μm. White arrows indicate colocalized sense and antisense 5’ETS RNAs. Green lines within insets indicated regions subjected to line-scan analysis (**D**) Line-scan analysis of relative fluorescence intensity across nucleolar cap regions (0–5 μm) showing colocalization of 5′ ETS sense (orange) and antisense (purple) RNA signals. (**E**) Model illustrating how rDNA double-strand breaks induce transcription of truncated, sense-antisense derived double-stranded rRNA species that are sensed by MDA5 and RIG-I, thereby activating innate immune signaling.

## Discussion

DNA Damage induced viral mimicry is mediated by both DNA and dsRNA pattern recognition (*52–54*). The source of immunogenic dsRNAs produced during DNA damage responses has remained enigmatic. We establish that rDNA double-strand breaks are a major site of immunogenic dsRNA generation in response to a wide spectrum of genotoxic agents. Breaks within the rDNA resulted in aberrant bidirectional rRNA transcription, generating overlapping sense and antisense rRNA species that rapidly engage cytosolic dsRNA-sensing pathways, MDA5 and RIG-I (**Fig. 5E**). ATM and ATR dependent rDNA silencing serves to limit both sense and antisense transcription emanating from broken rDNA (**Fig. 4E, F**), highlighting connections between these central DNA damage response regulators, rRNA transcription and processing, and innate immune activation.

Several groups have reported antisense RNA production from DNA double-strand breaks (*55–57*). While antisense transcription has not been observed at some genomic locations outside of the rDNA (*58*), direct comparisons of nuclease breaks showed that antisense transcripts from DSBs in the rDNA were more readily detected than from breaks in other genomic locations (*57*). RNA Pol II synthesizes antisense transcripts within intergenic regions between rDNA repeats in the absence of DNA damaging insults. This is thought to prevent readthrough transcription by RNA Pol I into intergenic rDNA regions in a manner that is necessary for ribosome biogenesis (*59*). The rDNA exists within a nucleolar condensate that contains high local concentrations of both RNA Pol I and Pol II. These features may be responsible for enhanced antisense transcription from rDNA break sites. It will also be of interest to determine how immunogenic sense-antisense derived rRNA rapidly enters the cytoplasm to become associated with RIG-I and MDA5 (**Fig. 2**). In regard to this issue, paired antisense-sense dsRNAs are reported to be preferentially exported from the nucleus (*60*).

RIG-I and MDA5 filaments have different dsRNA ligand specificities. MDA5 recognizes long dsRNAs while RIG-I activation requires 5’-triphosphorylated dsRNAs that minimally contain 10-20 base pairs of homology (*5, 61, 62*). Isolation of the active MDA5 or RIG-I filament had not been achieved, preventing direct identification of their cognate dsRNA ligands. We overcome this limitation by leveraging TRIM65 to purify the endogenous MDA5 filament following rDNA damage. This methodologic advance demonstrates complementary sense-antisense transcripts present within the vicinity of the 28S I-PpoI break site. Single molecule FISH studies further reveal directionality of antisense transcription along the length of the rDNA locus. Our efforts to purify endogenous RIG-I filaments in association with its endogenous dsRNA ligands using recombinant RIPLET, its cognate E3 ubiquitin ligase, have been thus far unsuccessful. RIG-I ligands could arise from unprocessed 5’-triphosphorylated dsRNAs at the 5’ETS start site (*63*), in accordance with the observed disruption in processing of immature rRNAs following rDNA damage (**Fig. 3**). Given the directionality of antisense production from rDNA damage, all sgRNAs employed in this study generated antisense transcripts within the 5’ETS, while Cas9 breaks within the 5’ETS generated antisense transcripts only within the 5’ETS. Interestingly, we observed the most robust ISG responses from breaks within the upstream 5’ETS sites (**Fig. 1G**), raising the possibility that dsRNAs emanating from this region generate active RIG-I ligands.

The nucleolus is proposed as an integrator of cellular stress responses (*47, 64*). Spontaneous DNA double-strand breaks frequently occur in rDNA, and rDNA damage associated transcriptional silencing is associated with hematopoietic stem cell senescence (*65, 66*). A ubiquitous feature of genotoxic chemo- and radiotherapies is to silence rRNA transcription and/or interfere with rRNA processing and packaging into the large and small subunits of the ribosome (*42*). Here we show that ATM and ATR activities promote rDNA silencing to a wide variety of genotoxic insults commensurate with their ability to mitigate dsRNA pattern recognition. This may explain findings that ATM and ATR inhibition can direct ISG responses to become primarily reliant on dsRNA pattern recognition (*21, 22*). While it is currently unclear how rDNA damage induced dsRNA pattern recognition influences therapeutic response, the efficacy of radio- and chemo- therapies relies on intact immune responses in syngeneic mouse models (*67*). Moreover, ATR inhibitors have been shown to enhance combined responses to DNA damaging and immune therapies (*68–70*). It will therefore be important to determine how rDNA derived dsRNAs arise in these settings and the underlying mechanisms that direct their engagement with MDA5 and RIG-I.

## Material and Methods

### Cell lines

The human telomerase reverse transcriptase (hTERT)-immortalized retinal pigment epithelial cell line RPE-1 was obtained from the American Type Culture Collection (ATCC). hTERT RPE-1 *p53* knockout (KO) cells were generated using plasmid-based CRISPR/Cas9-mediated knockout and used as the parental cell line to generate inducible expression of the site-specific homing endonuclease I-PpoI or AsiSI, or constitutive expression of Cas9. pLVX-pTUNER-C/ERT2-I-PpoI and pLVX-pTUNER-N/ERT2-AsiSI plasmids were used to generate I-PpoI and AsiSI lentiviruses, respectively. DLD1 cells expressing inducible I-PpoI or inducible AsiSI, and HEK293T cells were also used in this study. All cell lines were cultured in Gibco Dulbecco’s modified Eagle medium (DMEM, Thermo Fisher, 10564011) with 10% bovine calf serum (Cytiva, #SH30073.03HI) and 1% penicillin-streptomycin (Thermo Fisher Scientific, #15140122). Cells were maintained at 37 °C in a humidified incubator with 5% CO_2._ All cell lines were routinely tested and confirmed to be negative for mycoplasma using the MycoAlert Plus Mycoplasma Detection Kit (Lonza, #LT07-701).

### Plasmid-based CRISPR/Cas9-mediated knockout

hTERT RPE-1 *p53* KO cell line was generated by transfecting pSpCas9(BB)-2A-GFP(PX458) vector targeting *p53* into hTERT RPE-1 cell line (ATCC). 2 days post transfection, cells were sorted using a FACSAria sorter (BD) and then seeded at one cell per well in 96-well plate, and single clones were identified and marked under microscope 4 days later. Single clones were expanded, passaged, and validated by western blot. Single guide RNA (sgRNA) oligos used for *p53* knockout are: For #1, 5’-CACCG ATCCA CTCAC AGTTT CCAT-3’; Rev #1. 5’-aaacA TGGAA ACTGT GAGTG GATC-3’; For #2, 5’-CACCG AGCAC ATGAC GGAGG TTGTG-3’; Rev #2, 5’-aaacC ACAAC CTCCG TCATG TGCTC-3’.

### Lentiviral CRISPR/Cas9-mediated knockout

Lentiviruses for LentiCas9-mCherry (gift from Dr Junwei Shi, University of Pennsylvania), LentiCas9-blast, LentiCas9-Puro, and LentiGuide-Puro (Addgene, #52962, #104994, #52963) were produced and concentrated as previously described (*71*). Cells were infected with lentiviruses overnight in the presence of 8 μg/ml polybrene. Cells infected with LentiCas9-Blast were selected with 10 μg/ml blasticidin (Invivogen) until all uninfected control cells were eliminated. Cells infected with LentiCas9-Puro were selected with 20 μg/ml puromycin (Corning) with daily passages until all uninfected control cells were eliminated. Cells infected with LentiCas9-mCherry were sorted using an FACSAria sorter (BD). Knockouts were created by infecting Cas9-expressing cells with lentivirus produced using lentiGuide-Puro. To screen single colonies, cells were seeded one cell per well in 96-well plate and single clones were identified and marked under microscope 4 days later. Single clones were expanded, passaged, and validated by western blot. Otherwise, cells were cultured for 4 days before experimental treatment. Knockout efficiency was validated by immunoblotting using the corresponding antibodies. sgRNAs targeting HPRT, MDA5, RIG-I, LIG4, and XRCC4 were generated by inserting annealed duplex oligonucleotides into pLentiGuide-Puro at the BsmBI restriction site.

Oligos were obtained from Integrated DNA Technologies (IDT). sgRNA sequences used for gene knockout were as follows: sgHPRT #1, 5′-CACCG TTAGC TCCCA GGGAT CACAT-3′; sgHPRT #2, 5′-CACCG TCTGG GTCTG GATAA AAAGA-3′; sgHPRT #3, 5′-CACCG ATGGT TGTGA GCCAC CATG-3′; sgLIG4 #1, 5′-CACCG CAGAC AAAAG AGGTG AAGGG-3′; sgLIG4 #3-2, 5′-CACCG AACAG TTTGT GAAGT TTGTG-3′; sgXRCC4 #1, 5′-CACCG TACTG GGTTC AGAAA CAAGG-3′; sgXRCC4 #2, 5′-CACCG TTACT GATGG TCATT CAGCA-3′; sgXRCC4 #3, 5′-CACCG AAAAG GACAT CAAAC AAGA-3′.; sgControl: For #1,5’-CACCG CCCCG CCGCC CTCCC CTCC-3’; Rev #1, 5’-aaacG GAGGG GAGGG CGGCG GGGC-3’; For #2, 5’-CACCG CCAGT TGCTC TGGGG GAACA-3’; Rev #2. 5’-aaacT GTTCC CCCAG AGCAA CTGGC-3’; For #3. 5’- CACCG TCAGC AAAGG ACGAA ACAAA-3’; Rev #3, 5’-aaacT TTGTT TCGTC CTTTG CTGAC-3’. sgMDA5: For #1, 5’-CACCG GAGCA ATATA CTAGG ACTG-3’; Rev #1, 5’- aaacC AGTCC TAGTA TATTG CTCC-3’; For #2, 5’-CACCG CTCAG GGTTC ATGTA GCGGG-3’; Rev #2, 5’- aaacC CCGCT ACATG AACCC TGAGC-3’; For #3, 5’-CACCG CTGTT AGTCC CAGTA TCTG-3’; Rev #3, 5’-aaacC AGATA CTGGG ACTAA CAGC-3’. sgRIG-I: For #1, 5’- CACCG CTAGG GCATC CAAAA AGCCA-3’; Rev #1, 5’-aaacT GGCTT TTTGG ATGCC CTAGC-3’; For #2, 5’-CACCG AGATC AGAAA TGATA TCGGT-3’; Rev #2, 5’- aaacA CCGAT ATCAT TTCTG ATCTC-3’.

### Induction of rDNA breaks by I-PpoI and CRISPR/Cas9

Ribosomal DNA (rDNA) double-strand breaks were induced using two complementary approaches: inducible expression of the I-PpoI endonuclease and transfection of sgRNAs targeting the rDNA repeat locus in Cas9-expressing cells. For I-PpoI-mediated rDNA break induction, cells were treated with Shield-1 (1 μM, Aobious, #AON1848) and 4-hydroxytamoxifen (4-OHT; 2 μM, Sigma Aldrich, #H7904) for 48 h unless otherwise indicated. For CRISPR/Cas9-mediated rDNA break induction, single guide RNAs (sgRNAs) targeting corresponding rDNA were transfected into hTERT RPE-1 *p53* KO cells expressing Cas9, using Lipofectamine RNAimax (Thermo Fisher, #13778075) or Avalanche®-Omni transfection reagent (EZ Biosystems, #EZT-OMNI-1) according to manufacturer’s instructions. sgRNAs targeting the 5′- external transcribed spacer (5′-ETS), 18S, internal transcribed spacer 2 (ITS2), and 28S regions of rDNA were obtained from integrated DNA technologies (IDT-DNA). As a genome-wide DNA damage control, DSBs were induced using AsiSI endonuclease under the Shield-1 and 4-OHT induction conditions as described above.

The rDNA-targeting sgRNAs used in this study were as follows: Alt-R™ CRISPR-Cas9 hypoxanthine-guanine phosphoribosyltransferase (HPRT) control sgRNA, (crRNA targeting HPRT, IDT Oligos #1072533, paired with Cas9 tracrRNA); CD.Cas9.LJJV4159.AF (5′-ETS): AlTR1/rGr ArArGrUr CrArArCr CrCrArCr ArCrArCr GrArCrGr UrUrUrUr ArGrArGr CrUrArUr GrCrU /AlTR2; CD.Cas9.VRTB8269.AA (18S): AlTR1/rAr GrUrCrGr UrArArCr ArArGrGr UrUrUrCr CrGrUrGr UrUrUrUr ArGrArGr CrUrArUr GrCrU/AlTR2; CD.Cas9.PJPV0393.AS (ITS2): AlTR1/rCr GrGrGrCr ArCrGrGr GrArCrCr UrUrCrCr ArCrCrGr UrUrUrUr ArGrArGr CrUrArUr GrCrU/AlTR2; CD.Cas9.RFBD8173.AA (28S): AlTR1/rGr ArGrUrAr GrUrGrGr UrArUrUr UrCrArCr CrGrGrGr UrUrUrUr ArGrArGr CrUrArUr GrCrU/AlTR2; CD.Cas9.MXQV5533.AC(28S-1) /AltR1/rGrA rGrUrA rArCrU rArUrG rArCrU rCrUrC rUrUrA rGrUrU rUrUrA rGrArG rCrUrA rUrGrC rU/AltR2/; CD.Cas9.MXQV5533.AH(28S-2) /AltR1/rUrA rArUrU rArGrA rUrGrA rCrGrA rGrGrC rArUrU rGrUrU rUrUrA rGrArG rCrUrA rUrGrC rU/AltR2/ (IDT Oligos).

### Chemicals and Irradiation

Cells were seeded to allow sufficient space for proliferation, reaching a density of approximately 50% confluency at the time of collection. ATM i, ATR i or DNA-Pk i were administered 1 h prior to rDNA break induction. Drugs were maintained in the culture medium throughout the experiment, and the medium with or without drugs was changed every 2 days after treatment unless otherwise stated. Chemical inhibitors were used at the following concentrations: 5-ethynyl Uridine (EU) (1 mM, #AA00FBA1) (AA Blocks); ATM inhibitor (0.5 μM, AZD0156, #S8375), ATR inhibitor (0.25 μM, VE822), DNA-PK inhibitor (5 μM, NU7441 (KU-57788), S1092), Camptothecin (CPT) (500 nM, #S1288), Etoposide (20 μM, #S1225) (Selleck Chemicals); BMH-21 (2.5 μM, #HY-12484) (MedChem Express). For irradiation experiments, cells were seeded at ∼50% confluency at the time of treatment. Cells were irradiated using a ^137^Cs Gammacell irradiator (Nordion) with 0.76 Gy min^-1^. Inhibitors were administered 1 h prior to irradiation and maintained until collection, unless otherwise stated. The medium with or without inhibitors was refreshed every 2 days after treatment.

### Immunostaining

Cells were grown on 8-well chamber slides (Sycamore life sciences, #229168) or seeded onto coverslips in 24-well plates one day prior to treatment. Cells were fixed with pre-chilled 3% (w/v) paraformaldehyde (Sigma-Aldrich, #158127) for 15 min at room temperature, permeabilized with 0.5% Triton X-100 in 1x PBS for 10 min on ice, and blocked with blocking buffer (10% calf serum in 1x PBS) for 1 h at room temperature. For dsRNA staining, cells were fixed with methanol for 10 min at −20 °C and then staining was performed at 4 °C. Primary antibodies were diluted in blocking buffer and incubated overnight at 4 °C. Corresponding secondary antibodies were incubated for 1 h at room temperature. Slides were washed and mounted using VECTASHIELD mounting medium with DAPI (Vector Laboratories, #H-1200-10). Images were acquired using NIS-Elements software (Nikon) with a QImaging RETIGA-SRV camera connected to a Nikon Eclipse 80i microscope using a 40 x or 100 x objective. The following primary antibodies were used for immunostaining: anti-RPA194 (sc-48385), anti-UBF (sc-13125) (Santa Cruz Biotechnology); anti-fibrillarin (FBL) (ab5821) (Abcam); anti-γH2AX (05–636) (Millipore); anti-dsRNA (9D5, Ab00458-23.0) (Absolute antibody). The following secondary antibodies were used for immunofluorescence: Alexa Fluor™ 488- or 568-conjugated goat anti-rabbit or anti-mouse secondary antibodies (Invitrogen). Images were quantified using CellProfier 4.2.5 image analysis software and graphs were generated in GraphPad Prism 10.

### Western blotting

Western blotting was performed using standard methods. Briefly, cells were collected by trypsinization, washed with PBS, and lysed in NETN buffer (150 mM NaCl, 1% IGEPAL CA-630 (NP-40 alternative), 50 mM Tris-HCl, pH 7.4) with Turbo nuclease (Accelagen, #N0103M) in the presence of 1 mM MgCl_2_, cOmplete protease inhibitor cocktail (Sigma Aldrich, #5056489001) and phosphatase inhibitor cocktail (5 mM sodium fluoride, 1 mM sodium orthovanadate, 1 mM sodium pyrophosphate decahydrate, 1 mM β-glycerophophate) on ice for 45 min. Lysates were clarified by centrifugation at 13,000 rpm for 10 min at 4 °C, and supernatants were collected. Proteins were quantified using Pierce BCA Protein Assay Kit (Thermo 23225). Equal amount of protein was boiled with 4x NuPAGE LDS Sample Buffer (Thermo Scientific, NP0007) containing 10% β-mercaptoethanol at 94 °C for 5 minutes, then separated on 4%–12% Bis-Tris gels (Invitrogen) using MOPS buffer (Thermo Fisher Scientific, #NP000102) and transferred to 0.2 μm Amersham Protran nitrocellulose membranes (Cytiva, #10600004) at 350 mA for 2 h in ice-cold transfer buffer (20% methanol, 191 mM glycine, 25 mM Tris-base, 0.1% SDS, pH 8.3). Following Ponceau S staining (0.5% (w/v) Ponceau S in 25% acetic acid), membranes were blocked with 3% milk in TBST (0.1% Tween 20 in TBST) for 10 minutes and incubated with primary antibodies overnight at 4 °C. Membranes were washed with TBST and incubated with HRP-conjugated secondary antibodies (anti-mouse, NA931 and anti-rabbit, NA934, 1:25,000 dilution) (Amersham ECL HRP) for 1 h at room temperature. Blots were developed using Immobilon Forte Western HRP substrate (Millipore Sigma). The following primary antibodies were used for immunoblotting: anti-IFIH1 (MDA5) (Rabbit, #21775-1) (Proteintech); anti-RIG-I (Rabbit, #3743S), anti-phospho-STAT1 (Tyr701) (Rabbit, #9167S), anti-IFIT1 (ISG56) (Rabbit, #14769S), anti-ATM (Rabbit, #2843S), anti-GAPDH (Rabbit, #2118S), anti-HA tag (Rabbit, #3724S) (Cell Signaling Technology); anti-phospho-ATM (Rabbit, #ab81292) (Abcam); anti-phospho-IRF3 (S386) (Rabbit, #AP0995) (Abclonal); anti-γH2AX (05–636) (Millipore).

### TRIM65 purification

HEK293T cells were seeded in six 15 cm dishes at 12 million cells per dish one day before transfection. One hour before transfection, the medium was replaced with 25 ml of fresh medium. For each dish, 30 μg of pcDNA3-V5-SBP-HA-TRIM65 was transfected into cells using 42 μl Avalanche-Omni transfection reagent. Cells were cultured for 3 days with daily medium replacement. All subsequent procedures were performed in an RNase-free environment at 4 °C unless otherwise indicated. Cells from each dish were washed twice with 7 ml PBS, scraped into 4 ml PBS, pooled, and centrifuged to pellet. The cell pellet (∼2 ml) was resuspended in 18 ml modified hypotonic buffer (10 mM Tris-HCl pH 7.5, 10 mM KCl, 1.5 mM MgCl_2_, 5% glycerol, fresh 1 mM TCEP, and protease inhibitor cocktail), then aliquoted into two 50 ml LoBind tubes (Eppendorf, #0030122240) with 9 ml each and incubated for 15 min. Cells were lysed by gently passing the resuspension through a 25G needle fitted to a 10 ml syringe 7 times, avoiding bubbles.

The cell lysates were aliquoted into 1.7 ml tubes and sequentially centrifuged at 1,000 x g, 5,000 x g and 18,000 x g for 5 min each. The resulting cytoplasmic supernatant (∼10 ml) was carefully pooled into a 50 ml LoBind Eppendorf tube, mixed with an equal volume (10 ml) of modified RBB buffer (20 mM HEPES pH 7.5, 1.5 mM MgCl_2_, 1 M NaCl, 5% glycerol, 0.02% NP-40, fresh 1 mM TCEP, and protease inhibitor cocktail), and pre-cleared with 120 μl of TrueBlot Protein G magnetic beads (Rockland, #PG00-18-2) by gentle rotation for 15 min.

A 20 μl aliquot was taken for input, and the remaining cytoplasmic lysate was incubated with 240 μl Dynabeads MyOne Streptavidin C1 beads (Thermo Scientific, #65801D) for 45 min with gentle rotation. Beads were collected using a magnetic stand and gently resuspended in 1 ml modified RBB wash buffer (20 mM HEPES pH 7.5, 1.5 mM MgCl_2_, 500 mM NaCl, 5 % glycerol, 0.01 % NP-40, fresh 1 mM TCEP, and protease inhibitor cocktail) before being transferred to a 5 ml LoBind tube. Beads were washed twice with 4 ml modified RBB wash buffer for 15 min each with rotation, using a new 5 ml LoBind tube for each wash. Pull-down beads were resuspended in 1 ml modified RBB wash buffer containing 2 μl TurboNuclease (Accelagen, #N0103M) and transferred to 1.5 ml LoBind tube for overnight incubation at 4 °C. The next morning, beads were washed twice with 1 ml modified RBB wash buffer by gentle pipetting and resuspended in 240 μl of modified RBB wash buffer for TRIM65 pull-down on the same day.

### TRIM65 pull-down

Please also see reference (*76*) for experimental details. All experiments were performed in an RNase-free environment at 4 °C or on ice unless otherwise indicated. Three 15 cm dishes of RPE-1 cells were used for each sample. For each dish, cells were first washed twice with 7 ml PBS and once with 4 ml hypotonic buffer (10 mM Tris-HCl pH 7.5, 10 mM KCl, 1.5 mM MgCl_2_) containing protease inhibitor cocktail (Roche, #11873580001). Cells were scraped in the residual 0.7-1 ml hypotonic buffer, transferred to a 1.7 ml Eppendorf tube, and gently resuspended with a 1 ml pipette, followed by 15 min incubation. A 1 ml syringe with a 25G needle was used to gently pass the hypotonic buffer-resuspended cells through 7 times to lyse cells, avoiding bubbles. NP-40 was added to the cell lysates to a final concentration of 0.01 %, followed by sequential centrifuge at 1,000 x g, 5,000 x g and 18,000 x g for 5 min each.

The supernatant containing cytoplasmic lysate from three dishes was carefully transferred and pooled into a 5 ml LoBind Eppendorf tube (Eppendorf, #0030108302) to achieve ∼1.5 ml, then mixed with an equal volume (1.5 ml) of RBB buffer (20 mM HEPES pH 7.5, 1.5 mM MgCl_2_, 150 mM NaCl) with 0.01% NP-40 containing protease inhibitor cocktail. The mixture was precleared with TrueBlot Protein G magnetic beads (Rockland, #PG00-18-2) by gentle rotation for 15 min. A 100 μl aliquot was taken for input, and the remaining cytoplasmic lysate was incubated with 30 μl TRIM65-conjugated magnetic beads for 45 min with gentle rotation. Beads were collected using a magnetic stand and gently resuspended in 1 ml RBB buffer with 0.01% NP-40 before transferring to a 1.5 ml LoBind EP tube (Eppendorf, #0030108442). Beads were washed twice with 1 ml RBB buffer with 0.01% NP-40 for 15 min each with rotation, using a new 1.5 ml LoBind tube for each wash.

Pull-down beads were resuspended in 1 ml low-salt RBB buffer (20 mM HEPES pH 7.5, 1.5 mM MgCl_2_, 50 mM NaCl) with 0.01% NP-40. RNase A (Qiagen, #19101) was added to 1 ml resuspended pull-down beads and 100 μl input cytoplasmic lysate to a final concentration of 1 μg μl^-1^, followed by incubation at 4°C for 15 min. RNase A-digested beads were washed twice with 1 ml RBB buffer with 0.01% NP-40 by gentle pipetting and then resuspended in 300 μl RBB buffer with 0.01% NP-40. Aliquots of 20 μl and 280 μl were taken for pull-down western blot analysis and RNA-seq, respectively.

For western blot, 20 μl cytoplasmic lysate and 20 μl beads were boiled with 7 μl 4x NuPAGE LDS Sample Buffer (Thermo Scientific, #NP0007) containing 10% β-mercaptoethanol at 95°C for 5 minutes. For RNA-seq, an equal volume of 2x DNA/RNA shield (Zymo) was added to both the input lysate (50 μl) and the beads suspension (280 μl), followed by pipetting to mix. Proteinase K (Thermo Scientific, #25530049) was added to a final concentration of 1 mg ml^-1^ and incubated at room temperature for 30 min. An equal volume of DNA/RNA lysis buffer was then added to the input lysate-DNA/RNA Shield mixture (100 μl) and the beads suspension-DNA/RNA Shield mixture (560 μl), mixed by pipetting, and incubated at room temperature for 10 min. Beads were collected using a magnetic stand, and the supernatant was saved for RNA purification.

### RNA purification for TRIM65 pull-down

RNA for RNA-seq library preparation was purified using the Quick DNA/RNA Kit (Zymo, #D7005) according to the manufacturer’s instructions. Briefly, input RNA and eluted TRIM65 pull-down RNA that had been mixed with DNA/RNA shield and DNA/RNA lysis buffer were combined with an equal volume of ethanol and passed through a Zymo-Spin IC column by centrifugation. The column was washed with DNA/RNA wash buffer, followed by on-column DNase I digestion for 15 min at room temperature. After digestion, the column was washed once with DNA/RNA prep buffer and twice with DNA/RNA wash buffer before elution with 20 μl of 65 °C nuclease-free water. RNA concentration was initially measured by a NanoDrop spectrophotometer and diluted accordingly for validation on an Agilent High Sensitivity RNA ScreenTape using a TapeStation 4150 (Agilent Technologies).

### RNA-seq library preparation

RNA-seq libraries were prepared using the SMARTer® Stranded Total RNA-Seq Kit v3 - Pico Input Mammalian (Takara Bio, #634486) and SMARTer® RNA Unique Dual Index Kit – 96U Set A (Takara Bio, #634452) according to the manufacturer’s instructions. In brief, 10 ng of input RNA and all pull-down RNA samples (since the total RNA amount was less than 10 ng) were used for library preparation. For non-RNase A treated samples, fragmentation was performed at 94°C for 4 min, and for RNase A digested samples, fragmentation was omitted before first-strand cDNA synthesis. Illumina adapters with barcodes were added through PCR amplification: 5 cycles for non-RNase A treated samples and 10 cycles for RNase A treated samples, followed by ribosomal cDNA depletion. The final cDNA libraries were amplified by 13 PCR cycles for non-RNase A treated samples and 16 PCR cycles for RNase A treated samples, then purified using NucleoMag beads (Takara, #744970.5). cDNA library concentrations were initially measured by NanoDrop and diluted accordingly for validation on an Agilent High Sensitivity D1000 ScreenTape using an TapeStation 4150 (Agilent Technologies). cDNA libraries were normalized to a concentration of 6 nM prior to pooling and pooled in equimolar amounts. The library pools were clustered and sequenced on a NovaSeq X platform with 150-bp paired-end reads, using sequencing services provided by Innomics.

### Analysis of TRIM65-pulldown data

TRIM65-pulldown peaks were called with JAMM (version 1.0.7rev6, parameters: -d y -t paired - b 50 -w 1 -m normal) (*72*) using two replicates of the respective pull-down as foreground and the two replicates of the corresponding input condition as background. The peaks called in the knockout of MDA5 were treated as non-specific background signals and subtracted from the peaks called in the presence of MDA5.

To determine the enrichment of repetitive elements in pulldown samples compared to input controls, we employed a previously established mapping-free, graph-based sequence clustering method (*73, 74*). Briefly, we sampled 1,000,000 reads from the TRIM65-pulldown profiling datasets. These reads were processed through a clustering pipeline (*75*) that conducts all-to-all pairwise comparisons to construct a sequence graph. In this graph, reads serve as vertices (nodes), and edge weights reflect the sequence similarity scores between them. The graph was subsequently partitioned into distinct clusters representing connected communities. Next, we utilized RepeatMasker to annotate the reads within each cluster for the presence of various repeat families. Finally, the largest clusters (those with the highest read counts) for each sample were visualized to evaluate the composition of repetitive elements.

### Northern blotting

RNA was purified using the Quick DNA/RNA Kit (Zymo, #D7005) according to the manufacturer’s instructions. Briefly, cells were resuspended in 1x DNA/RNA Shield and Proteinase K (Thermo Scientific, #25530049) was added to a final concentration of 1 mg ml^-1^ and incubated at room temperature for 30 min. An equal volume of DNA/RNA lysis buffer was then added and mixed by pipetting, and incubated at room temperature for 10 min. The mixture was combined with an equal volume of ethanol and passed through a Zymo-Spin IC column by centrifugation. The column was washed with DNA/RNA wash buffer, followed by on-column DNase I digestion for 15 min at room temperature. After digestion, the column was washed once with DNA/RNA prep buffer and twice with DNA/RNA wash buffer before elution with 20 μl of 65°C nuclease-free water. RNA concentration was initially measured by a NanoDrop spectrophotometer. 0.375 μg of total RNA was resolved on 1% formaldehyde agarose gels, transferred overnight onto a nylon Hybond-N+ membrane (Cytiva) by capillary transfer, UV-crosslinked (120 mJ/cm^2^), and hybridized with 1-5 nM biotin-labelled probes. Signals were detected using 1:300 diluted streptavidin-HRP (Thermo 89880). Northern blot probes are as follows: ITS2 probe: 5’-CGCAC CCCGA GGAGC CCGGA GGCAC CCCCG G-3’; 5’ETS probe: 5’-CACGA ACGTC CGCCC CTCGC CCGTC GCGGC TCGG-3’.

### Click chemistry for labeling nascent RNA

For detecting nascent RNA synthesis with the EU-Click reaction, cells were labeled with 1 mM 5′-ethynyl uridine (EU) (AA Blocks, #AA00FBA1) for 30 min before harvesting and processed using the Click-iT® Plus Alexa Fluor 488 Picolyl Azide kit (Thermo Fisher Scientific, #C10641) according to the manufacturer’s instructions. In brief, cells were fixed with 3 % PFA for 15 min, permeabilized with 0.5 % Triton X-100 for 10 min, and subjected to the click reaction.

For EU-Biotin click reactions, cells were labeled with 1 mM EU (AA Blocks, #AA00FBA1) for 1 h before harvesting and fixed with pre-chilled 3% PFA for 15 min at room temperature, permeabilized with 0.5% Triton X-100 in PBS for 10 min at room temperature. Cells were washed with 1x PBS twice for 5 min each. After washing, the click reaction cocktail (2 mM CuSO_4_, 20 μM Biotin-PEG4-Picolyl azide (BroadPharm, #BP-22954) and 100 μM sodium L-ascorbate (Sigma Aldrich, #A4034) in 1X PBS) was added to each chamber then incubated at room temperature for 1 h. Subsequent steps were carried out with the PLA method described below, using either mouse or rabbit anti-biotin antibodies in conjunction with the antibody for the protein of interest.

### Proximity Ligation Assay (PLA)

After 5′-ethynyl uridine (EU) incubation for 1 h, cells were fixed with pre-chilled 3 % paraformaldehyde (PFA) for 15 min at room temperature, and incubated in CSK buffer (10 mM PIPES, 100 mM NaCl, 300 mM sucrose, 3 mM MgCl2, 1 mM EGTA, and 0.5% Triton X-100) for 10 min at 4 °C, and subjected to the biotin click reaction for 1 h at room temperature. After the click reaction, cells were blocked with 10% fetal bovine serum in PBS for 1 h at room temperature and incubated with primary antibodies overnight at 4°C. The next day, after washing with 1x PBS twice, cells were incubated with pre-mixed Duolink® *In situ* PLA® probes, anti-Rabbit MINUS (Sigma, #DUO92005), and anti-Mouse PLUS (Sigma, #DUO92001) probes for 1 h at 37°C. Subsequent steps in PLA were carried out using the Duolink® *In situ* Detection Reagents Red (Sigma, #DUO92008) according to the manufacturer’s instructions. Slides were washed and mounted using VECTASHIELD mounting medium with DAPI (Vector Laboratiroies, #H-1200-10). Images were acquired with a QImaging RETIGA-SRV camera connected to a Nikon Eclipse 80i microscope at 40× objective. The following primary antibodies were used for PLA: anti-Biotin (Mouse, #SAB4200680) (Sigma); anti-Biotin (Rabbit, #5597S), anti-RIG-I (Rabbit, #3743S) (Cell Signaling Technology); anti-MDA5 (IFIH1) (Rabbit, #21775-1) (Proteintech).

### Single-molecule RNA fluorescence in situ hybridization (smFISH)

RNA smFISH probes were designed using the Stellaris Probe Designer (Biosearch Technologies), synthesized by Integrated DNA Technologies (IDT-DNA), and prepared in-house. The assay was performed using Stellaris RNA FISH Buffers (Biosearch Technologies) according to the manufacturer’s instructions. Briefly, cells grown on glass coverslips were fixed in 3% PFA for 10 minutes at room temperature, permeabilized in 70% ethanol for at least 1 hour at 4 °C, and hybridized with 75 nM strand-specific Stellaris probe sets targeting the indicated rRNA regions in Hybridization buffer (Biosearch Technologies) overnight at 37 °C. Following post-hybridization washes twice with Buffer A (Biosearch Technologies), nuclei were counterstained with DAPI, and samples were mounted using Dako Fluorescence Mounting Medium for imaging (Agilent). Images were acquired using a Nikon fluorescence microscope, and smFISH signals were quantified using imageJ.

smFISH probes are as follows:

**5′ETS probes: Sense:** S#1, 5′-TCAGA GGACC CCGCC GGC-3′; S#2, 5′-AGCGA GGGCT GTCTG CCG-3′; S#3, 5′-CGACA ACCAC TGGAG GCG-3′; S#4, 5′-GAGGG GGGCC GCCCG CAA-3′; S#5, 5′-ACGGC ACCCC CACCG CCG-3′; S#6, 5′-GCGCA AACCC CCCGA GAG-3′; S#7, 5′-CCCAG GCGGA GCCGA CGC-3′; S#8, 5′-CTCCA GGAGC ACCGC AAG-3′; S#9, 5′-CTGAG GGACA ACCCG GAG-3′; S#10, 5′-CACCG TTCGG CCTCG GGC-3′; S#11, 5′-CGGGG GCGGG AACGA CAC-3′; S#12, 5′-CACGC GCGGA CACCG CGG-3′; S#13, 5′-CGACG AGCTC CCTCA GGA-3′; S#14, 5′-AAACC GCCTC GAACC CCA-3′; S#15, 5′-CGTCT CGTCT CGTCT CAC-3′; S#16, 5′-TTCCC CGCGT GGGAG GGG-3′; S#17, 5′-GAGCA CGGGA CGTGC GCT-3′; S#18, 5′-CGCGC GCACC CGCCA GAG-3′; S#19, 5′-ACCGC GATCG CTCAC ACG-3′; S#20, 5′-CGTCA CACCG GCCCG AAC-3′.

**Antisense** AS#1, 5′-CTCTG ACGCG GCAGA CAG-3′; AS#2, 5′-AGTGG TTGTC GACTT GCG-3′; AS#3, 5′-CGGCC CCCCT CCGCG GCG-3′; AS#4, 5′-CCGTC GTGCT GCCCT CTC-3′; AS#5, 5′-GGGGT TTGCG CGAGC GTC-3′; AS#6, 5′-CTCCG CCTGG GCCCT TGC-3′; AS#7, 5′-TGCTC CTGGA GCGCT CCG-3′; AS#8, 5′-GAACG GTGGT GTGTC GTT-3′; AS#9, 5′-CTCCG GTCGC CGCCG CGG-3′; AS#10, 5′-TGAGG GAGCT CGTCG GTG-3′; AS#11, 5′-GGTTC GAGGC GGTTT GAG-3′; AS#12, 5′-GAGAC GAGAC GCGCC CCT-3′; AS#13, 5′-CACGC GGGGA AGGGC GCC-3′; AS#14, 5′-CCTGC TCTCG GTGAG CGC-3′; AS#15, 5′-GTCCC GTGCT CCCCT CTG-3′; AS#16, 5′-GGGTG CGCGC GGGCC GTG-3′; AS#17, 5′-AGCGA TCGCG GTGGG TTC-3′; AS#18, 5′-GCCGG TGTGA CGCGT GCG-3′; AS#19, 5′-GGCCG GCCGC CGAGG GGC-3′; AS#20, 5′-CCGTT CTGCC TCCGA CCG-3′.

18S probes: Sense S#1, 5′-GTCAA TTCCT TTAAG TTT-3′; S#2, 5′-ACTCC TGGTG GTGCC CTT-3′; S#3, 5′-TGAGG TTTCC CGTGT TGA-3′; S#4, 5′-CTGTC CGTGT CCGGG CCG-3′; S#5, 5′-GAGCT ATCAA TCTGT CAA-3′; S#6, 5′-CACCC ACGGA ATCGA GAA-3′; S#7, 5′-TAAGA ACGGC CATGC ACC-3′; S#8, 5′-AGACA AATCG CTCCA CCA-3′; S#9, 5′-GTTCG TTATC GGAAT TAA-3′; S#10, 5′-AGTTA GCATG CCAGA GTC-3′; S#11, 5′-CGCTC GGGGG TCGCG TAA-3′; S#12, 5′-AAGAA GTTGG GGGAC GCC-3′; S#13, 5′-TGGCT GAACG CCACT TGT-3′; S#14, 5′-CAGAC CTGTT ATTGC TCA-3′; S#15, 5′-CCGGA CATCT AAGGG CAT-3′; S#16, 5′-GAGCC AGTCA GTGTA GCG-3′; S#17, 5′-GCGTA GGGTA GGCAC ACG-3′; S#18, 5′-TGGGG TTCAA CGGGT TAC-3′; S#19, 5′-AATTG CAATC CCCGA TCC-3′; S#20, 5′-ATTCC TCGTT CATGG GGA-3′; S#21, 5′-GCGGT GTGTA CAAAG GGC-3′; S#22, 5′-ACCAT CCAAT CGGTA GTA-3′; S#23, 5′-CGATC CGAGG GCCTC ACT-3′; S#24, 5′-CAGGG CCGTG GGCCG ACC-3′.

**Antisense** AS#1, 5′-AAAGT CTTTG GGTTC CGG-3′; AS#2, 5′-GGAGT ATGGT TGCAA AGC-3′; AS#3, 5′-TTGAC GGAAG GGCAC CAC-3′; AS#4, 5′-AGCCT GCGGC TTAAT TTG-3′; AS#5, 5′-ACACG GGAAA CCTCA CCC-3′; AS#6, 5′-CCGGA CACGG ACAGG ATT-3′; AS#7, 5′-AGCTC TTTCT CGATT CCG-3′; AS#8, 5′-CATGG CCGTT CTTAG TTG-3′; AS#9, 5′-GGAGC GATTT GTCTG GTT-3′; AS#10, 5′-ACGAA CGAGA CTCTG GCA-3′; AS#11, 5′-AACTA GTTAC GCGAC CCC-3′; AS#12, 5′-GTCGG CGTCC CCCAA CTT-3′; AS#13, 5′-TAGAG GGACA AGTGG CGT-3′; AS#14, 5′-AGCCA CCCGA GATTG AGC-3′; AS#15, 5′-TAACA GGTCT GTGAT GCC-3′; AS#16, 5′-TAGAT GTCCG GGGCT GCA-3′; AS#17, 5′-CGCGC TACAC TGACT GGC-3′; AS#18, 5′-AGCGT GTGCC TACCC TAC-3′; AS#19, 5′-CGCGG GTAAC CCGTT GAA-3′; AS#20, 5′-CCATT CGTGA TGGGG ATC-3′; AS#21, 5′-GGATT GCAAT TATTC CCC-3′; AS#22, 5′-GAACG AGGAA TTCCC AGT-3′; AS#23, 5′-GTGCG GGTCA TAAGC TTG-3′; AS#24, 5′-TTGAT TAAGT CCCTG CCC-3′; AS#25, 5′-GTCGC TACTA CCGAT TGG-3′; AS#26, 5′-TTAGT GAGGC CCTCG GAT-3′; AS#27, 5′-CCCAC GGCCC TGGCG GAG-3′; AS#28, 5′-CTGAG AAGAC GGTCG AAC-3′; AS#29, 5′-GACTA TCTAG AGGAA GTA-3′.

**ITS2 probes: Sense** S#1, 5′-TTTCA CACCA CGGGG AGG-3′; S#2, 5′-GGGAC CGGAC TCCGG AGA-3′; S#3, 5′-GGCCA GACGA GACAG CAA-3′; S#4, 5′-AGAGG GGGTT GCCTC AGG-3′; S#5, 5′-CCCCC CCCCC GCCCA AGA-3′; S#6, 5′-TTCCT GGCGC GGCAC GTC-3′; S#7, 5′-GACGC ACCGG GAGGA GGC-3′; S#8, 5′-TTGGC GAGGG CGCTC CCG-3′; S#9, 5′-AAGAG TCGTA CGAGG TCG-3′; S#10, 5′-GATTG ATCGG CAAGC GAC-3′; S#11, 5′-CCGGA GGCAC CCCCG GGG-3′; S#12, 5′-AGCCG CGCAC CCCGA GGA-3′; S#13, 5′-GGCCC TGCGA GGGAA CCC-3′; S#14, 5′-GGGAC GGAGG GCCCC CGG-3′; S#15, 5′-CCGCC GGGTC TGCGC TTA-3′; S#16, 5′-CGGCA AGAGG AGGGC GGA-3′; S#17, 5′-AGGGG GAAGG GGCGG GCG-3′; S#18, 5′-CACGC AGGGC CCGCG GGG-3′; S#19, 5′-CCCGC CACCC GACGC GTG-3′; S#20, 5′-CGGGC GCGCC CCCCT CTC-3′; S#21, 5′-TCGCG GGCGG CGGCG GCG-3′; S#22, 5′-CTCTC TTTCC CTCTC CGT-3′; S#23, 5′-GAACT CGGCC CGAGC CGG-3′; S#24, 5′-ACCGC AGGCG GCGGC CAC-3′; S#25, 5′-CCCCC GAGGG AGGAA CCC-3′; S#26, 5′-CGCGC GCGGC GCGAG GGA-3′; S#27, 5′-ACGAA CCCCG AACCC CGA-3′.

**Antisense** AS#1, 5′-TCCGA CCCCT CTCCG GAG-3′; AS#2, 5′-CGGTC CCGTT TGCTG TCT-3′; AS#3, 5′-TCTGG CCGGC CTGAG GCA-3′; AS#4, 5′-CCCCT CTCCT CTTGG GCG-3′; AS#5, 5′-GGGGG GGGGG ACGTG CCG-3′; AS#6, 5′-CCAGG AAGGG CCTCC TCC-3′; AS#7, 5′-GTGCG TCGTC GGGAG CGC-3′; AS#8, 5′-CTCGC CAAAT CGACC TCG-3′; AS#9, 5′-CGACT CTTAG CGGTG GAT-3′; AS#10, 5′-GCTCG TGCGT CGATG AAG-3′; AS#11, 5′-GCAGC TAGCT GCGAG AAT-3′; AS#12, 5′-TGCAG GACAC ATTGA TCA-3′; AS#13, 5′-CTTCG AACGC ACTTG CGG-3′; AS#14, 5′-GTTCC TCCCG GGGCT ACG-3′; AS#15, 5′-TGTCT GAGCG TCGCT TGC-3′; AS#16, 5′-ATCAA TCGCC CCCGG GGG-3′; AS#17, 5′-CCTCC GGGCT CCTCG GGG-3′; AS#18, 5′-CGCGG CTGGG GGTTC CCT-3′; AS#19, 5′-CAGGG CCCGC CGGGG GCC-3′; AS#20, 5′-CCGTC CCCCT AAGCG CAG-3′; AS#21, 5′-CCGGC GGCGT CCGCC CTC-3′; AS#22, 5′-TTCCC CCTCC CCCCG CGG-3′; AS#23, 5′-CTGCG TGGTC ACGCG TCG-3′; AS#24, 5′-TGGCG GGGGG GAGAG GGG-3′; AS#25, 5′-GCTGA GAGAG ACGGG GAG-3′; AS#26, 5′-CCCGC GAAGA CGGAG AGG-3′; AS#27, 5′-AAGAG AGAGC CGGCT CGG-3′; AS#28, 5′-CTGCG GTCCG GGTTC CTC-3′; AS#29, 5′-TCGGG GGGCT CCCTC GCG-3′; AS#30, 5′-CGCGC GGCTC GGGGT TCG-3′; AS#31, 5′-GTTCG TCGGC CCCGG CCG-3′.

**28S probes: Sense** S#1, 5′-GACGC CCGCC GCAGC TGG-3′; S#2, 5′-CGGGC TCCCC GGGGG CGG-3′; S#3, 5′-GGGGG GGCGC GCCGG CGC-3′; S#4, 5′-CGCGA CGAGA CGTGG GGT-3′; S#5, 5′-CCCCG CCCCC AGCGG ACG-3′; S#6, 5′-CGGGG GGGTA GGGCG GGG-3′; S#7, 5′-GGGAA CGGGG GGCGG ACG-3′; S#8, 5′-CGCCG CGCGC CGAGG AGG-3′; S#9, 5′-GGACC CGGCG GGGGG GAC-3′; S#10, 5′-TGCTA GGCGC CGGCC GAG-3′; S#11, 5′-TCCGC ACCAG TTCTA AGT-3′; S#12, 5′-TCGCG ATGCT TTGTT TTA-3′; S#13, 5′-CGTCA ACACC CGCCG CGG-3′; S#14, 5′-TTTCT TCACT TTGAC ATT-3′; S#15, 5′-CGTTT ACCCG CGCTT CAT-3′; S#16, 5′-AGAGA GTCAT AGTTA CTC-3′.

**Antisense** AS#1, 5′-GGATC GCCCC AGCTG CGG-3′; AS#2, 5′-CGTCG CGGCC GCCCC CGG-3′; AS#3, 5′-GCGCG CGCGT CCGCT GGG-3′; AS#4, 5′-GGGAG CGGTC GGGCG GCG-3′; AS#5, 5′-GGTTC GTCCC CCCGC CCT-3′; AS#6, 5′-CCCCG GCCCC GTCCG CCC-3′; AS#7, 5′-CCCCC GCCGG GTCCG CCC-3′; AS#8, 5′-GGCCG CGGTT CCGCG CGG-3′; AS#9, 5′-TCGCC TCGGC CGGCG CCT-3′; AS#10, 5′-GCCGA CTTAG AACTG GTG-3′; AS#11, 5′-CAGGG GAATC CGACT GTT-3′; AS#12, 5′-CAAAG CATCG CGAAG GCC-3′; AS#13, 5′-GGGTG TTGAC GCGAT GTG-3′; AS#14, 5′-CCCAG TGCTC TGAAT GTC-3′; AS#15, 5′-ATTCA ATGAA GCGCG GGT-3′; AS#16, 5′-GGCGG GAGTA ACTAT GAC-3′.

## Statistics and reproducibility

All statistical analyses were performed using GraphPad Prism 10 software. All data were assembled into figures using Adobe Illustrator 2025. Significance was calculated using an unpaired two-tailed Student’s *t*-test. *p* values are shown in the figures. All source data and brief instructions for data organization are provided. Bar graphs showing mean ± SEM of independent experimental results, with the number of replicates indicated in the figure legends. Microscopy related images were randomly acquired. Western blots (WB) checking protein expression level after overexpression and/or CRISPR knockout were performed at least three times. **** p < 0.0001, *** p < 0.001, ** p < 0.01, * p < 0.05, and n.s.: not significant. No statistical methods were used to predetermine sample size.

## Acknowledgments

We thank members of the Greenberg, Li, and Liu laboratories for critical discussions. Research reported in this publication was supported by the National Cancer Institute of the National Institutes of Health. The content is solely the responsibility of the authors and does not necessarily represent the official views of the National Institutes of Health.

## Author contributions

Conceptualization: JC, SK, QL, KFL, RAG; Methodology: JC, SK, SC, ZM, YJ, KFL, QL, RAG; Investigation: JC, SK, QL, RAG; Visualization: JC, SK, SC, QL, RAG; Funding acquisition: RAG; Project administration: JC, SK, RAG; Supervision: RAG; Writing – original draft: JC, SK, RAG; Writing – review & editing: JC, SK, YJ, KFL, QL, RAG.

## Data, code, and materials availability

All data are available in the main text or the supplementary materials.

