## Supplemental Figures for "rDNA breaks activate dsRNA pattern recognition through sense-antisense transcription"

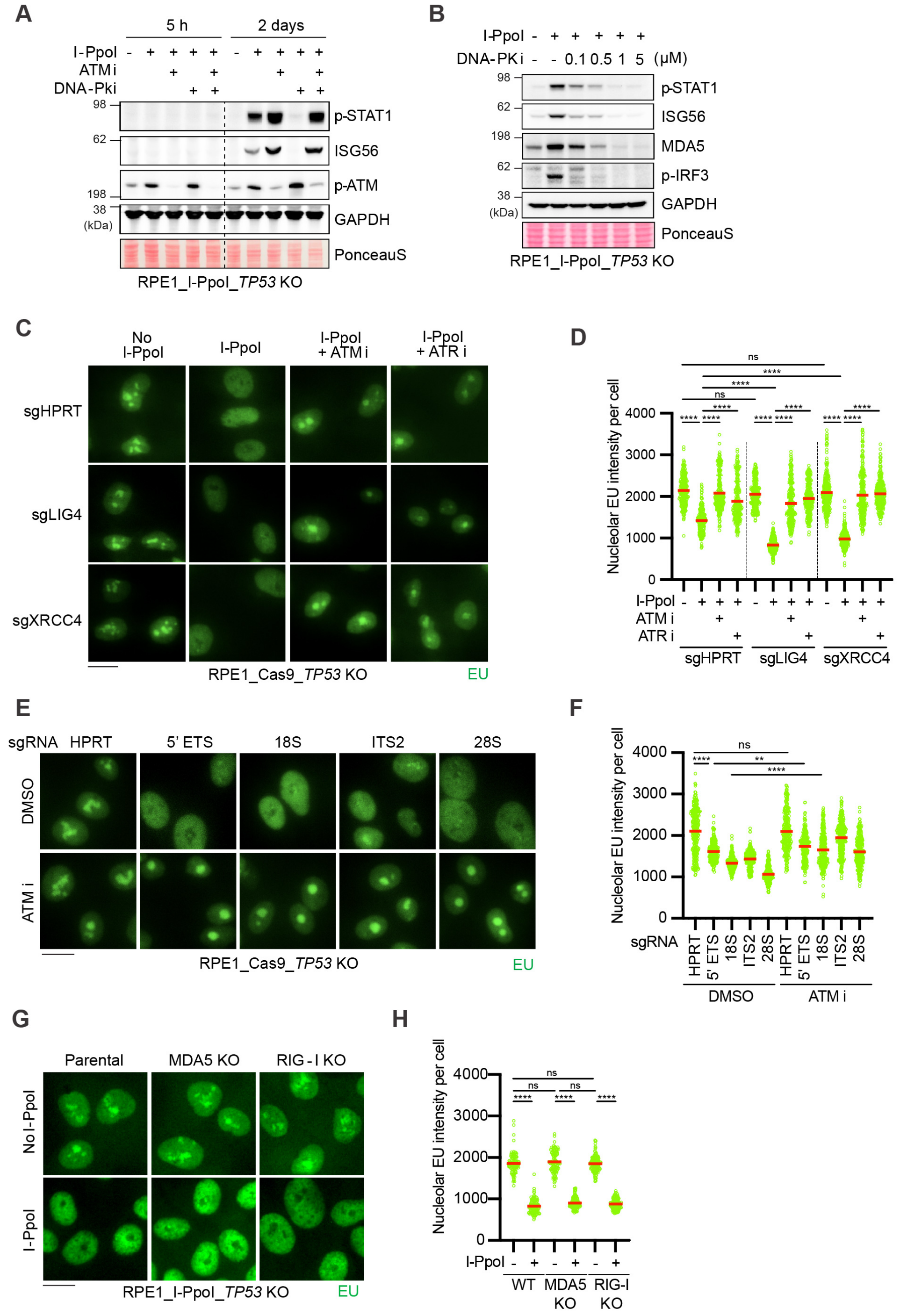
A
5 h
2 days
I-PpoI
ATM i
DNA-Pki
p-STAT1
ISG56
p-ATM
GAPDH
PonceauS
RPE1_I-PpoI_TP53 KO
B
I-PpoI
DNA-PK i
0.1 0.5 1 5 (μM)
p-STAT1
ISG56
MDA5
p-IRF3
GAPDH
PonceauS
RPE1_I-PpoI_TP53 KO
C
No I-PpoI
I-PpoI
I-PpoI + ATM i
I-PpoI + ATR i
sgHPRT
sgLIG4
sgXRCC4
RPE1_Cas9_TP53 KO
EU
D
Nucleolar EU intensity per cell
I-PpoI
ATM i
ATR i
sgHPRT
sgLIG4
sgXRCC4
E
sgRNA HPRT 5' ETS 18S ITS2 28S
DMSO
ATM i
RPE1_Cas9_TP53 KO
EU
F
Nucleolar EU intensity per cell
sgRNA
HPRT
5' ETS
18S
ITS2
28S
DMSO
ATM i
G
Parental
MDA5 KO
RIG - I KO
No I-PpoI
I-PpoI
RPE1_I-PpoI_TP53 KO
EU
H
Nucleolar EU intensity per cell
I-PpoI
WT
MDA5 KO
RIG-I KO

**Supplementary Figure 1. ATM and ATR kinase-dependent transcriptional silencing at rDNA breaks restricts innate immune responses.**

**(A, B)** Immunoblots of whole-cell lysates showing STAT1 phosphorylation and ISG56 expression following I-PpoI-mediated rDNA breaks in the presence of ATM or DNA-PK inhibition in I-PpoI-inducible hTERT RPE-1 I-Ppo1 *p53* knockout cells. **(C-H)** Immunofluorescence of nucleolar RNA synthesis visualized by EU-click labeling, with representative images and quantification shown. (**C, E, G**) Representative images of nucleolar nascent RNA synthesis. (**D, F, H**) Quantification of nucleolar EU intensity per cell. (**E, F**) rDNA-targeting sgRNAs are transfected into Cas9-expressing hTERT RPE-1 *p53* KO cells for 48 h. (**C-H**) At least 150 cells were quantified per experiment. Three independent experiments were performed, and one representative result is displayed. Scale bar, 20 μm. Statistical significance was determined by an unpaired two-tailed Student's *t*-test. **** $p < 0.0001$, ** $p < 0.01$, ns: not significant.

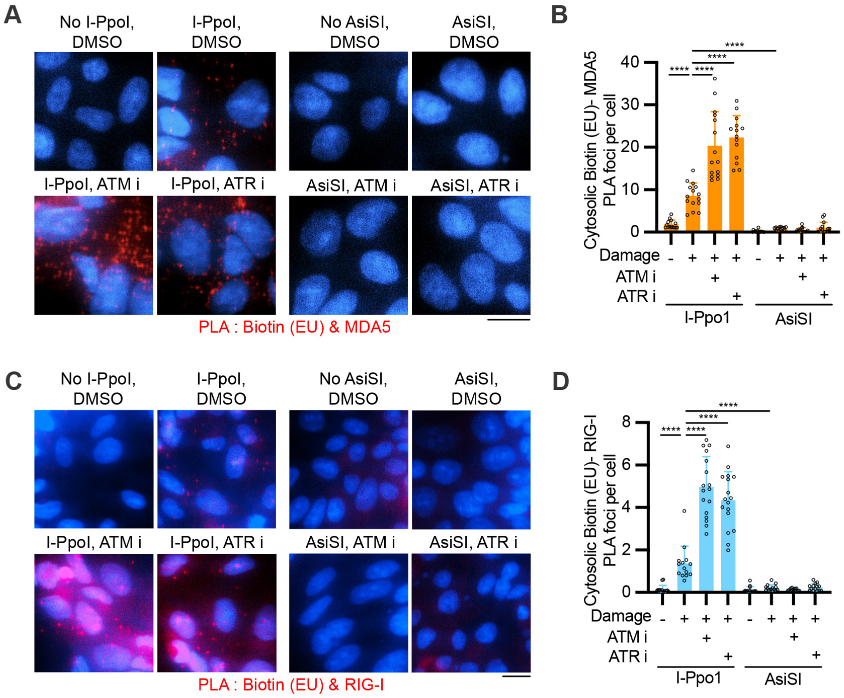
A
No I-PpoI, DMSO
I-PpoI, DMSO
No AsiSI, DMSO
AsiSI, DMSO
I-PpoI, ATM i
I-PpoI, ATR i
AsiSI, ATM i
AsiSI, ATR i
PLA : Biotin (EU) & MDA5
B
Cytosolic Biotin (EU)- MDA5 PLA foci per cell
40
30
20
10
0
****
Damage
ATM i
ATR i
I-Ppo1
AsiSI
C
No I-PpoI, DMSO
I-PpoI, DMSO
No AsiSI, DMSO
AsiSI, DMSO
I-PpoI, ATM i
I-PpoI, ATR i
AsiSI, ATM i
AsiSI, ATR i
PLA : Biotin (EU) & RIG-I
D
Cytosolic Biotin (EU)- RIG-I PLA foci per cell
8
6
4
2
0
****
Damage
ATM i
ATR i
I-Ppo1
AsiSI

**Supplementary Figure 2. I-PpoI-mediated rDNA breaks induce cytosolic nascent RNA sensing.**

(**A, C**) Representative images of proximity ligation assay (PLA) following EU labeling and biotin click reaction after I-PpoI- or AsiSI-mediated DNA damage. PLA signals indicate proximity between biotin-labeled nascent RNA (EU) and MDA5 or RIG-I. Scale bar, 20 μm. **(B, D)** Quantification of cytosolic PLA signals between EU-labeled nascent RNA and MDA5 (B) or RIG-I (D). (**A-D**) Experiments were performed in I-PpoI- or AsiSI-inducible DLD1 *p53* KO cells. Each dot represents the mean value from at least 50 cells. Three independent experiments were performed, and one representative result is displayed. Bars represent mean ± SEM. Statistical significance was determined by an unpaired two-tailed Student's *t*-test. **** $p < 0.0001$.

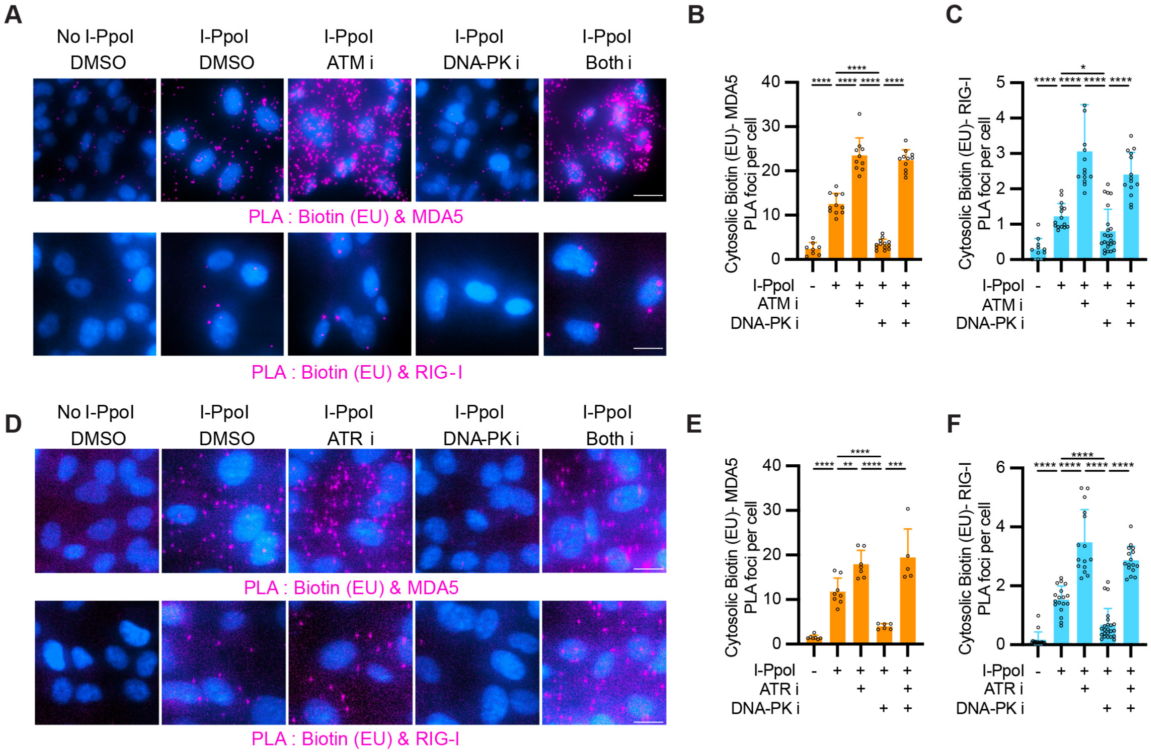
A
No I-PpoI DMSO
I-PpoI DMSO
I-PpoI ATM i
I-PpoI DNA-PK i
I-PpoI Both i
PLA : Biotin (EU) & MDA5
PLA : Biotin (EU) & RIG-I
B
Cytosolic Biotin (EU)- MDA5 PLA foci per cell
I-PpoI
ATM i
DNA-PK i
C
Cytosolic Biotin (EU)- RIG-I PLA foci per cell
D
No I-PpoI DMSO
I-PpoI DMSO
I-PpoI ATR i
I-PpoI DNA-PK i
I-PpoI Both i
PLA : Biotin (EU) & MDA5
PLA : Biotin (EU) & RIG-I
E
Cytosolic Biotin (EU)- MDA5 PLA foci per cell
ATR i
F
Cytosolic Biotin (EU)- RIG-I PLA foci per cell

**Supplementary Figure 3. DNA damage kinases regulate cytosolic RNA-sensing PLA signals upon rDNA breaks.**

(**A, D**) Representative images of proximity ligation assay (PLA) following EU labeling and biotin click chemistry after I-PpoI-mediated rDNA breaks. PLA signals indicate proximity between biotin-labeled nascent RNA (EU) and MDA5, or RIG-I. Scale bar, 20 µm. **(B, C, E, F)** Quantification of cytosolic PLA signals between EU-labeled nascent RNA and MDA5 (B, E), or RIG-I (C, F). (**A-F**) Experiments were performed in I-PpoI-inducible hTERT RPE-1 *p53* KO cells. Each dot represents the mean value from at least 50 cells. Bars represent mean ± SEM. Statistical significance was determined by an unpaired two-tailed Student's *t*-test. **** $p < 0.0001$, *** $p < 0.001$, ** $p < 0.01$, * $p < 0.05$.

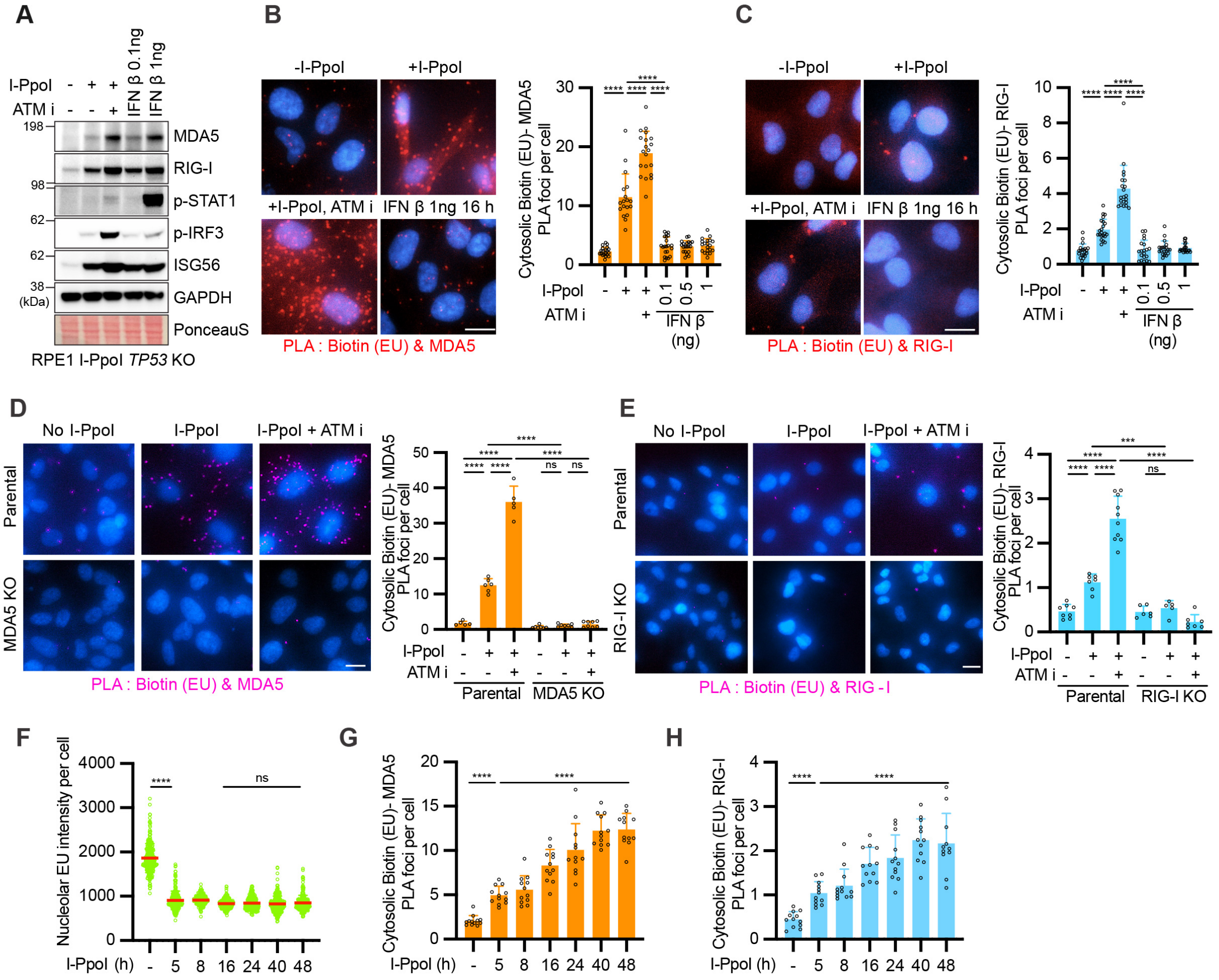
A
B
C
D
E
F
G
H
I-PpoI
ATM i
IFN β 0.1ng
IFN β 1ng
MDA5
RIG-I
p-STAT1
p-IRF3
ISG56
GAPDH
PonceauS
(kDa)
RPE1 I-PpoI TP53 KO
-I-PpoI
+I-PpoI
+I-PpoI, ATM i
IFN β 1ng 16 h
PLA : Biotin (EU) & MDA5
PLA : Biotin (EU) & RIG-I
Cytosolic Biotin (EU)- MDA5 PLA foci per cell
Cytosolic Biotin (EU)- RIG-I PLA foci per cell
IFN β (ng)
No I-PpoI
I-PpoI + ATM i
Parental
MDA5 KO
RIG-I KO
PLA : Biotin (EU) & RIG - I
Nucleolar EU intensity per cell
I-PpoI (h)
ns

**Supplementary Figure 4. Validation of rDNA break-specific cytosolic nascent RNA sensing PLA.**

(**A**) Immunoblots of whole-cell lysates showing STAT1 phosphorylation and ISG56 expression following I-PpoI-mediated rDNA breaks or interferon beta (IFN-β) treatment for 16 h in hTERT I-PpoI-inducible RPE-1 *p53* knockout cells. (**B, C, D, E**) Representative images of PLA following EU labeling and biotin click chemistry after I-PpoI-mediated rDNA breaks. PLA signals indicate proximity between biotin-labeled nascent RNA (EU) and MDA5 or RIG-I. Scale bar, 20 μm. Corresponding quantification of cytosolic PLA signals is shown on the right. (**F-H**) Time-course analysis of I-PpoI-mediated rDNA break induction, nucleolar EU intensity, and cytosolic PLA signals. At least 150 cells were quantified per experiment. For PLA analysis, each dot represents the mean value from at least 50 cells. Bars represent mean ± SEM. Statistical significance was determined by an unpaired two-tailed Student's *t*-test. **** $p < 0.0001$, *** $p < 0.001$, ns: not significant.

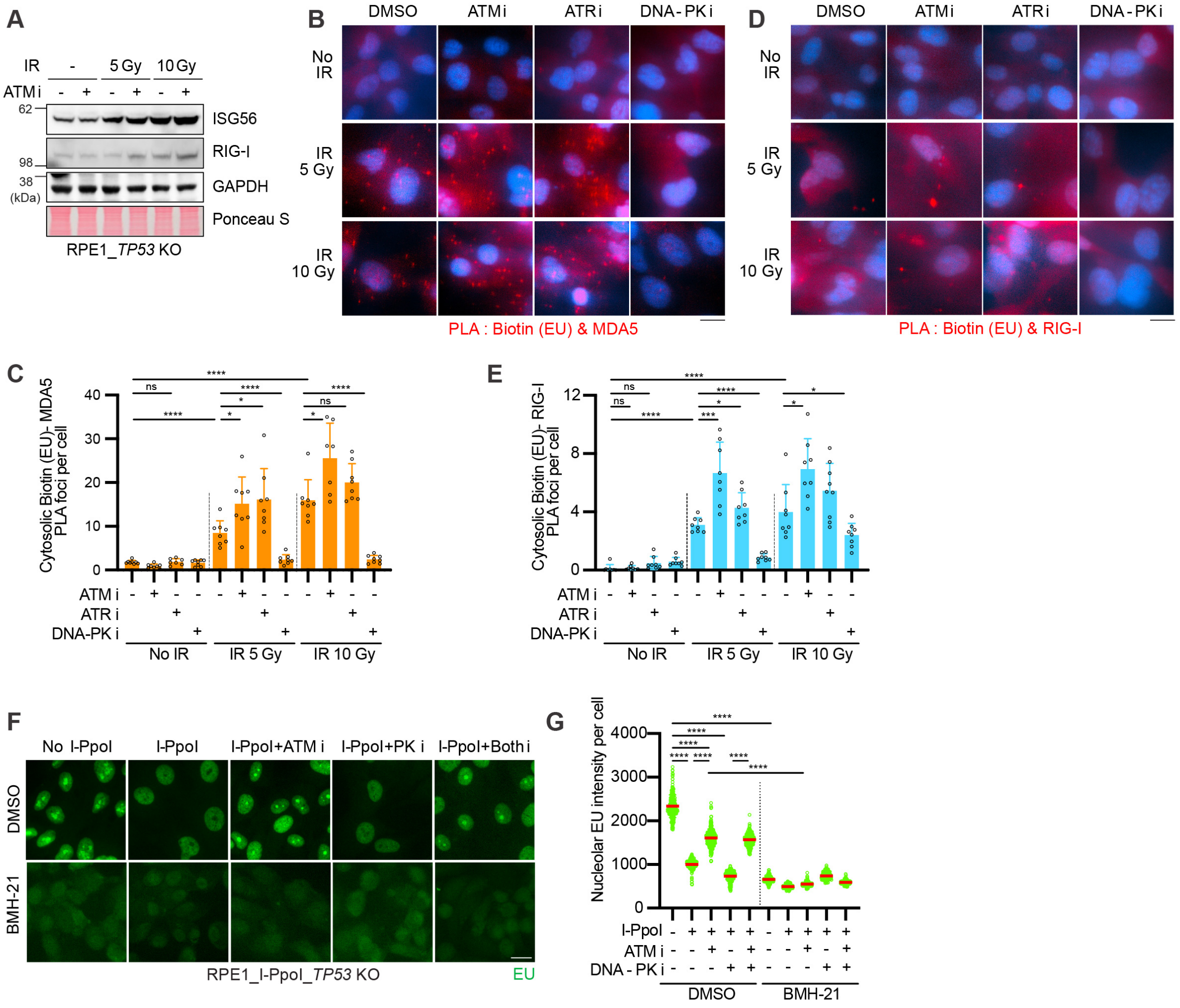
A
IR - 5 Gy 10 Gy
ATM i - + - + - +
62
ISG56
RIG-I
98
38
(kDa)
GAPDH
Ponceau S
RPE1_TP53 KO
B
DMSO ATM i ATR i DNA-PK i
No IR
IR 5 Gy
IR 10 Gy
PLA : Biotin (EU) & MDA5
D
DMSO ATM i ATR i DNA-PK i
No IR
IR 5 Gy
IR 10 Gy
PLA : Biotin (EU) & RIG-I
C
Cytosolic Biotin (EU)- MDA5 PLA foci per cell
ATM i
ATR i
DNA-PK i
No IR
IR 5 Gy
IR 10 Gy
E
Cytosolic Biotin (EU)- RIG-I PLA foci per cell
ATM i
ATR i
DNA-PK i
No IR
IR 5 Gy
IR 10 Gy
F
No I-PpoI
I-PpoI
I-PpoI+ATM i
I-PpoI+PK i
I-PpoI+Both i
DMSO
BMH-21
RPE1_I-PpoI_TP53 KO
EU
G
Nucleolar EU intensity per cell
I-PpoI
ATM i
DNA - PK i
DMSO
BMH-21

**Supplementary Figure 5. Irradiation (IR)-induced DNA breaks activate innate immune responses and cytosolic RNA sensing.**

(**A**) Immunoblots of whole-cell lysates showing STAT1 phosphorylation and ISG56 expression following irradiation (IR). (**B, D**) Representative images of proximity ligation assay (PLA) following EU labeling and biotin click chemistry after IR-induced DNA damage. PLA signals indicate proximity between biotin-labeled nascent RNA (EU) and MDA5 (B) or RIG-I (D). Scale bar, 20 μm. (**C, E**) Quantification of cytosolic PLA signals between EU-labeled nascent RNA and MDA5 (C) or RIG-I (E). For PLA analysis, each dot represents the mean value from at least 50 cells. (**F**) Representative images of nucleolar nascent RNA synthesis visualized by EU-click labeling following RNA polymerase I inhibition with BMH-21. (**G**) Quantification of nucleolar EU intensity. (**F, G**) At least 150 cells were quantified per experiment. Bars represent mean ± SEM. Statistical significance was determined by an unpaired two-tailed Student's *t*-test. **** $p < 0.0001$, *** $p < 0.001$, * $p < 0.05$, ns: not significant.

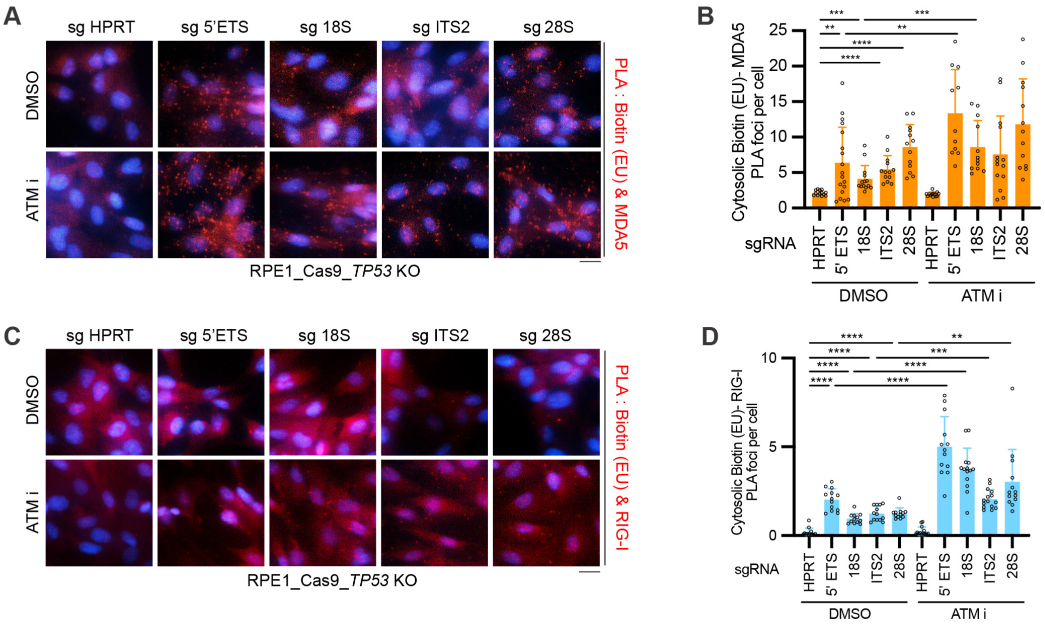
A
sg HPRT
sg 5'ETS
sg 18S
sg ITS2
sg 28S
DMSO
ATM i
PLA : Biotin (EU) & MDA5
RPE1_Cas9_TP53 KO
B
Cytosolic Biotin (EU)- MDA5 PLA foci per cell
sgRNA
HPRT
5' ETS
18S
ITS2
28S
DMSO
ATM i
C
PLA : Biotin (EU) & RIG-I
D
Cytosolic Biotin (EU)- RIG-I PLA foci per cell

**Supplementary Figure 6. Site-specific rDNA breaks induced by sgRNAs trigger cytosolic RNA-sensing PLA signals.**

(**A-D**) PLA analysis following Cas9-mediated rDNA break generation using sgRNAs targeting the 5'-ETS, 18S, ITS2, or 28S regions of rDNA repeats. (**A, C**) Representative images of PLA following EU labeling and biotin click chemistry after rDNA targeting with the indicated sgRNAs. Scale bar, 20 μm. (**B, D**) Quantification of cytosolic PLA signals between EU-labeled nascent RNA and MDA5 (B), or RIG-I (D). For PLA analysis, each dot represents the mean value from at least 50 cells. (**B, D**) Bars represent mean ± SEM. Statistical significance was determined by an unpaired two-tailed Student's *t*-test. **** $p < 0.0001$, *** $p < 0.001$, ** $p < 0.01$.

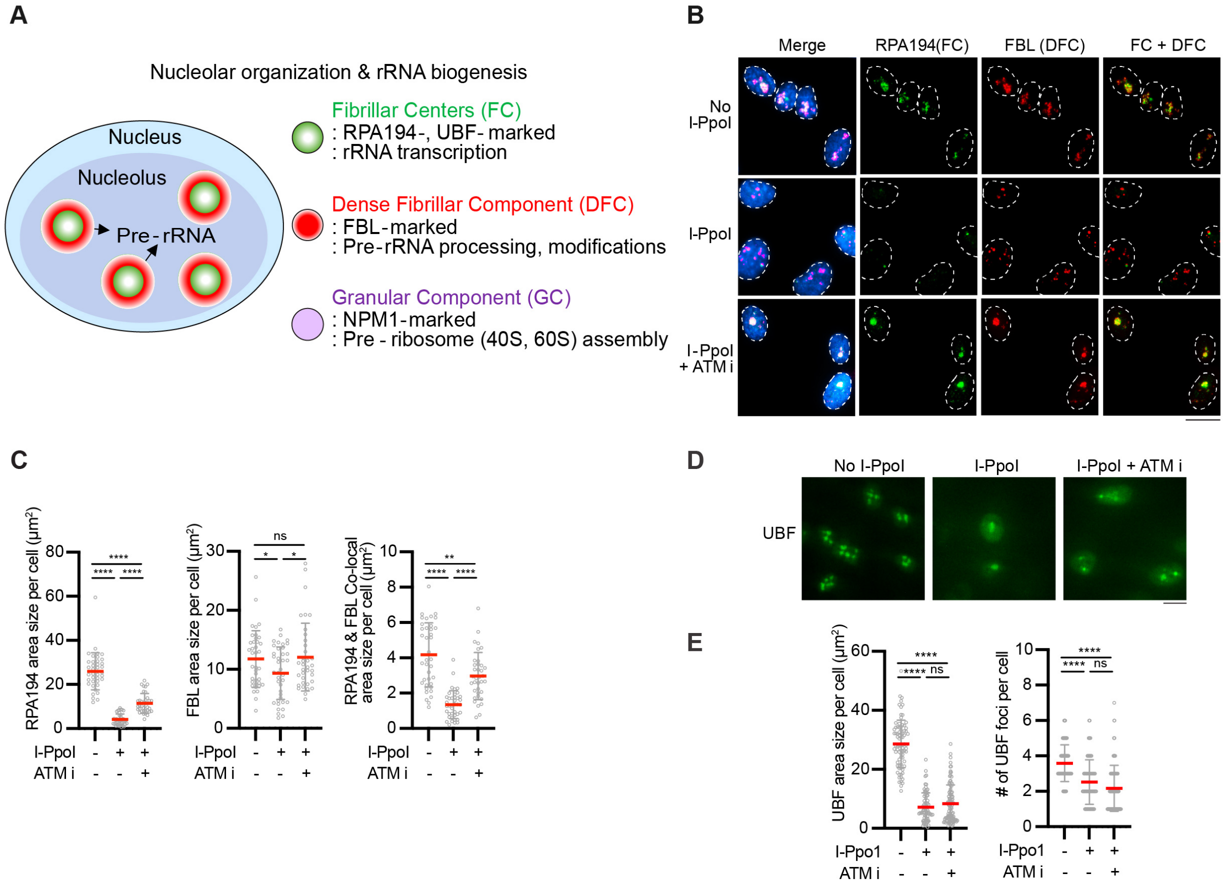
A
Nucleolar organization & rRNA biogenesis
Nucleus
Nucleolus
Pre-rRNA
Fibrillar Centers (FC)
: RPA194-, UBF-marked
: rRNA transcription
Dense Fibrillar Component (DFC)
: FBL-marked
: Pre-rRNA processing, modifications
Granular Component (GC)
: NPM1-marked
: Pre - ribosome (40S, 60S) assembly
B
Merge
RPA194(FC)
FBL (DFC)
FC + DFC
No I-PpoI
I-PpoI
I-PpoI + ATM i
C
RPA194 area size per cell (μm²)
FBL area size per cell (μm²)
RPA194 & FBL Co-local area size per cell (μm²)
I-PpoI
ATM i
****
ns
*
**
D
No I-PpoI
I-PpoI
I-PpoI + ATM i
UBF
E
UBF area size per cell (μm²)
### of UBF foci per cell
I-Ppo1
ATM i
****
ns

**Supplementary Figure 7. rDNA breaks disrupt nucleolar structure.**

(**A**) Schematic illustration of nucleolar organization and rRNA biogenesis. **(B)** Representative immunostaining images showing nucleolar subcompartments marked by RPA194, fibrillar center (FC) marker, and fibrillarin (FBL), a dense fibrillar component (DFC) marker, and their spatial colocalization. Scale bar, 20 μm. **(C)** Quantification of RPA194 area, FBL area, and co-localization area ($\mu m^2$) per cell. (**D**) Representative images of UBF immunostaining. Scale bar, 20 μm. (**E**) Quantification of UBF foci size and number per cell. Three independent experiments were performed, and one representative result is displayed. (**C, E**) At least 150 cells were quantified per experiment. Statistical significance was determined by an unpaired two-tailed Student's *t*-test. **** $p < 0.0001$, ** $p < 0.01$, * $p < 0.05$, ns: not significant.

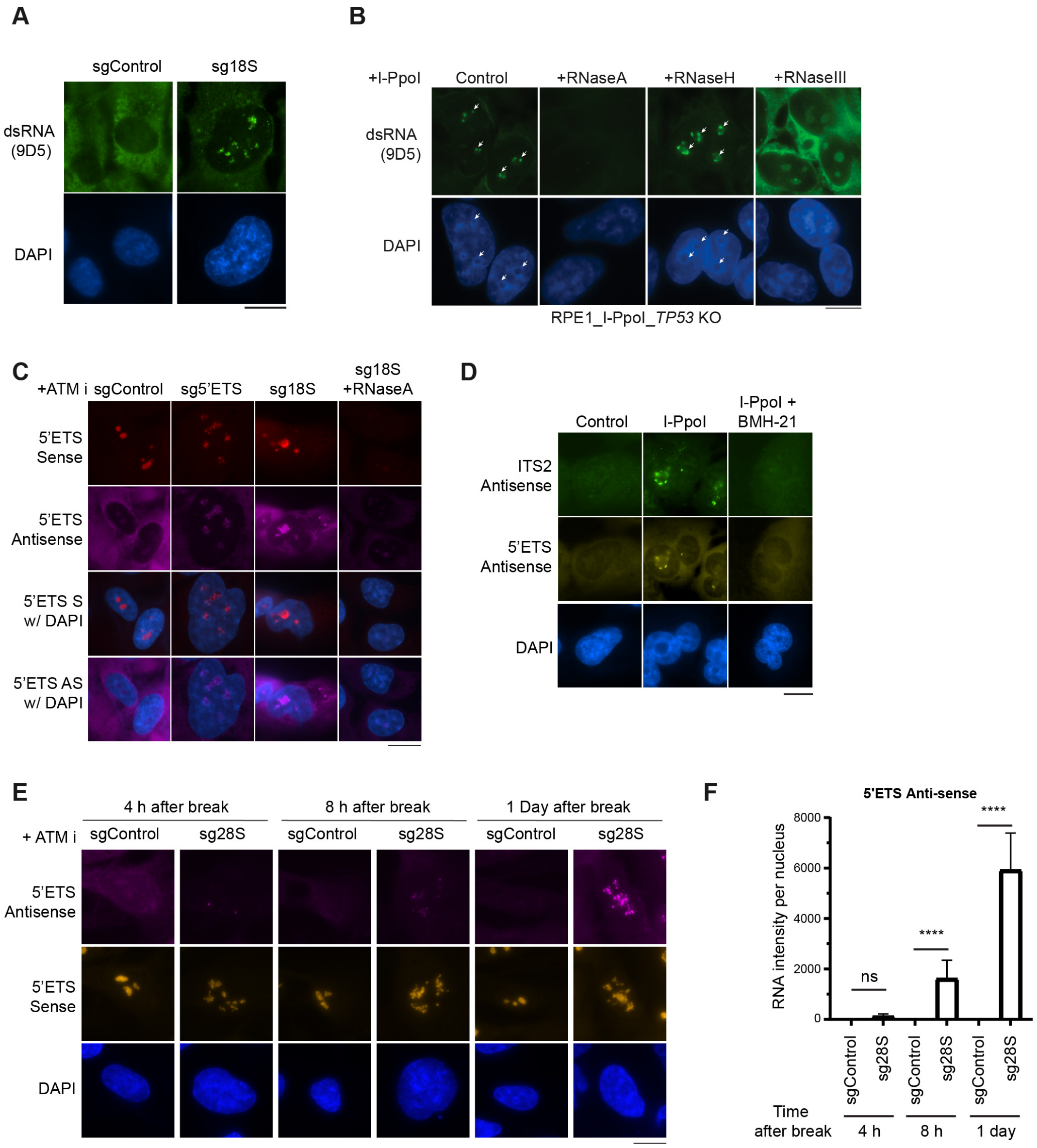

A
sgControl
sg18S
dsRNA (9D5)
DAPI
B
+I-PpoI
Control
+RNaseA
+RNaseH
+RNaseIII
dsRNA (9D5)
DAPI
RPE1_I-PpoI_TP53 KO
C
+ATM i
sgControl
sg5'ETS
sg18S
sg18S +RNaseA
5'ETS Sense
5'ETS Antisense
5'ETS S w/ DAPI
5'ETS AS w/ DAPI
D
Control
I-PpoI
I-PpoI + BMH-21
ITS2 Antisense
5'ETS Antisense
DAPI
E
4 h after break
8 h after break
1 Day after break
+ ATM i
sgControl
sg28S
5'ETS Antisense
5'ETS Sense
DAPI
F
5'ETS Anti-sense
RNA intensity per nucleus
8000
6000
4000
2000
0
ns
****
****
sgControl
sg28S
Time after break
4 h
8 h
1 day

**Supplementary Figure 8. Characterization of rDNA damage-induced sense and antisense rRNA transcription.**

**(A, B)** Representative dsRNA immunostaining images following sg18S- or I-PpoI-mediated break induction. (**B**) Corresponding RNases treatment before dsRNA immunostaining. (**C**) Representative strand-specific smFISH images of nascent RNA following gRNA-mediated targeting of the 5' ETS and 18S regions in the presence of ATM inhibition. RNase A treatment was included as a negative control. Sense and antisense transcripts were visualized using 5' ETS probes. (**D**) Representative strand-specific smFISH images of nascent RNA following I-PpoI-mediated rDNA break generation in the presence of the RNA polymerase I inhibitor BMH-21. 5' ETS and ITS2 antisense transcripts were visualized. (**E, F**) Time-course analysis of 5' ETS sense and antisense transcripts visualized by smFISH following gRNA-mediated break induction at the 28S rDNA region. (**F**) Quantification of 5' ETS antisense smFISH RNA intensity per nucleus. At least 50 cells were quantified per experiment. Bars represent mean ± SEM. Statistical significance was determined by an unpaired two-tailed Student's *t*-test. **** $p < 0.0001$, ns: not significant. (**A-E**) Scale bar, 20 μm.
